# Genome-wide dynamics of RNA synthesis, processing and degradation without RNA metabolic labeling

**DOI:** 10.1101/520155

**Authors:** Mattia Furlan, Eugenia Galeota, Nunzio Del Gaudio, Erik Dassi, Michele Caselle, Stefano de Pretis, Mattia Pelizzola

## Abstract

The kinetic rates of RNA synthesis, processing and degradation determine the dynamics of transcriptional regulation by governing both the abundance and the responsiveness to modulations of premature and mature RNA species. The study of RNA dynamics is largely based on the integrative analysis of total and nascent transcription, with the latter being quantified through RNA metabolic labeling. We describe here INSPEcT-, a computational method based on mathematical modeling of intronic and exonic expression, able to derive the dynamics of transcription from steady state or time course profiling of just total RNA, without requiring any information on nascent transcripts. Our approach closely recapitulates the kinetic rates obtained through RNA metabolic labeling, improves the ability to detect changes in transcripts half-lives, reduces the cost and complexity of the experiments, and can be adopted to study experimental conditions where nascent transcription cannot be readily profiled. Finally, we applied INSPEcT- to the characterization of post-transcriptional regulation landscapes in dozens of physiological and disease conditions. This approach was included in the INSPEcT Bioconductor package, which can now unveil RNA dynamics from steady state or time course data, with or without the profiling of nascent RNA.

## Main

Since the development of microarrays first, and high-throughput sequencing later on, the investigation of the transcriptional activity of genes has been mostly based on the quantification of total RNA (Mortazavi et al., 2008). While bringing about a revolution in the field of transcriptional regulation, the quantification of absolute and differential expression provides only a glimpse of the complexity of cellular gene expression programs. Indeed, abundance and responsiveness to modulations of premature and mature RNA species are set by the combined action of three key steps: premature RNA synthesis, processing of premature into mature RNA, and degradation of the latter (Orphanides and Reinberg, 2002). These steps are governed by corresponding kinetic rates, which collectively determine the RNA dynamics of transcripts (Fig. 1A). However, the specific contribution of each step of the RNA life cycle cannot be deconvoluted from an aggregate quantity like the amount of total RNA because, in principle, infinite combinations of kinetic rates can generate the same absolute expression level.

**Figure 1.**
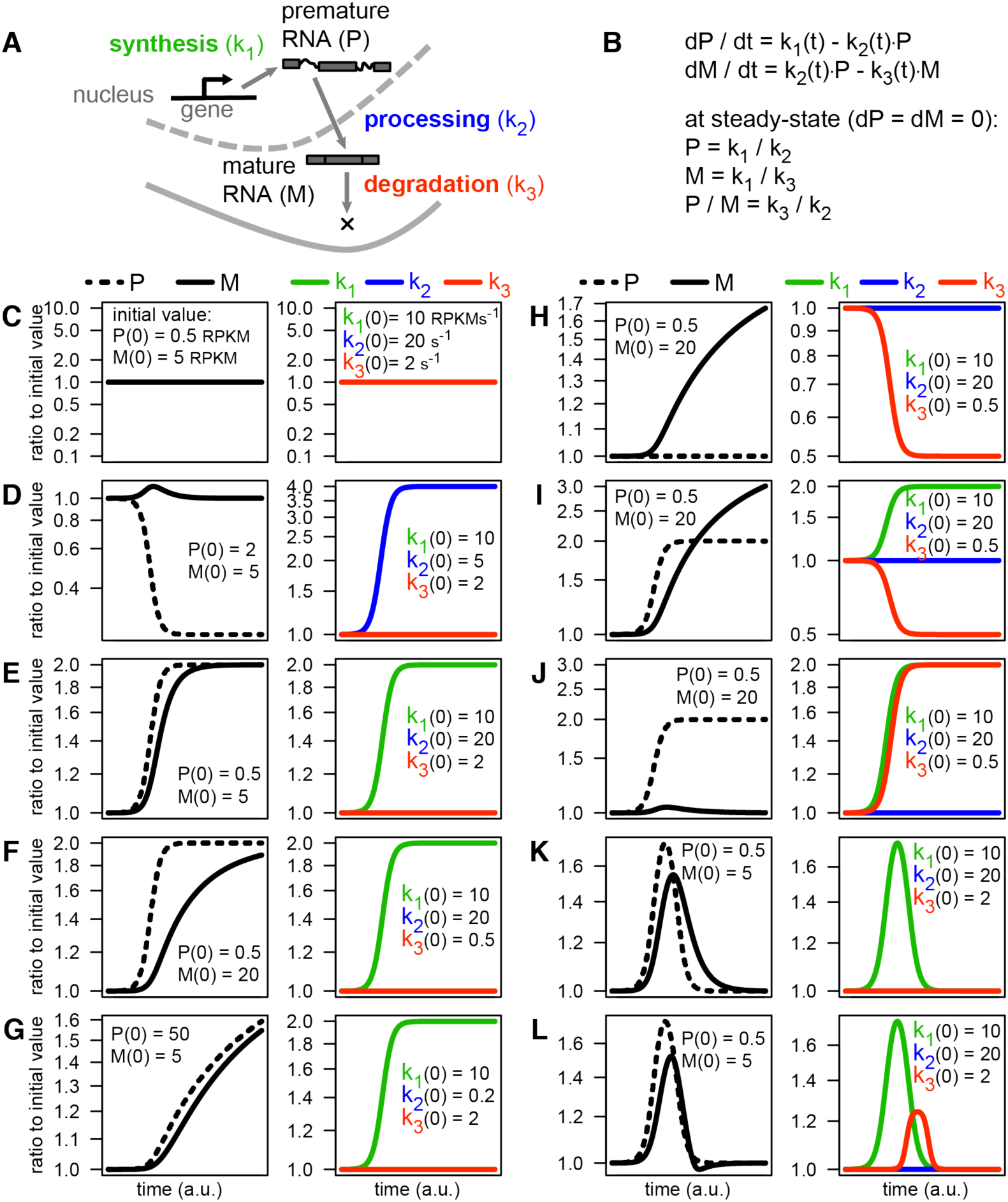
The influence of RNA kinetic rates on RNA abundance and responsiveness. (**A**) Schematic representation of the RNA life cycle, governed by the kinetics rates of synthesis, processing and degradation. (**B**) Deterministic mathematical model of the RNA life cycle based on Ordinary Differential Equations (ODEs), including the solution of the system at steady state. (**C-L**) Solutions of the ODE system following the modulation of the kinetic rates: each example reports, for premature and mature RNA species (left) and for the kinetic rates (right), the ratio to the initial time point. Initial values are indicated within each panel.

For decades the study of RNA dynamics relied solely on transcription blockage experiments. However, these methods allow the quantification of RNA half-lives only, are highly invasive, affect cell viability and could alter various pathways, RNA decay included (Wada and Becskei, 2017). To overcome these limitations, new methods have been developed that are based on the integrative analysis of total and nascent RNA. Nascent RNA can be metabolically labeled with biotinylated, 4-thiouridine modified (4sU) nucleotides, purified with streptavidin and then sequenced (Dolken et al., 2008; Miller et al., 2011; Rabani et al., 2011; Wissink et al., 2019). Alternatively, if the modified nucleotides are chemically derivatized prior to sequencing, reads from nascent transcripts can be *in silico* separated from pre-existing RNA (Baptista and Dölken, 2018; Herzog et al., 2017; Jürges et al., 2018; Riml et al., 2017; Schofield et al., 2018). These approaches have started to unveil how the modulation of RNA dynamics can determine gene-specific regulatory modes and elicit complex transcriptional responses (de Pretis et al., 2015; 2017; Furlan et al., 2019; Rabani et al., 2014).

Despite their advantages and popularity, methods based on RNA metabolic labeling are affected by various pitfalls, especially when limited amount of nascent RNAs is produced, and when aiming at studying very short responses (Baptista and Dölken, 2018). Moreover, these methods cannot be readily applied to model organisms, be it mammals (Matsushima et al., 2018) or plants (Sidaway-Lee et al., 2014), *in vivo*. For all these reasons, being able to study RNA dynamics from just total RNA would be a valuable alternative. A few studies have moved in this direction by using an integrative analysis of premature and mature RNA abundances (Gaidatzis et al., 2015; Gray et al., 2014; La Manno et al., 2018; Zeisel et al., 2011), yet they have fallen short of quantifying the full set of RNA kinetic rates and their modulation.

We describe here INSPEcT-, a computational approach that determines RNA dynamics from total RNA-seq data, thus avoiding all the downsides of RNA metabolic labeling. INSPEcT-quantifies the full set of kinetic rates from time course RNA-seq datasets, and enables the study of post-transcriptional regulation between steady state conditions. We used INSPEcT- to analyze different time-course RNA-seq datasets, ranging from conditions where gene expression programs are mostly controlled by transcriptional changes, to conditions where post-transcriptional regulation prevails. Finally, we used this method to characterize post-transcriptional regulation landscapes in dozens of tissue types and disease conditions. INSPEcT-is available within the INSPEcT Bioconductor package (http://bioconductor.org/packages/INSPEcT/), formerly developed by us for the analysis of RNA metabolic labeling data (de Pretis et al., 2015), and allows the user to study RNA dynamics on steady state or time-course data, with or without nascent RNA profiling.

## Results

### The quantification of RNA dynamics unveils the complexity of gene expression programs

At steady state, the abundance of premature RNA is equal to the ratio of its synthesis to its processing rate, and the quantity of its mature form is given by its synthesis to degradation rate ratio (Fig. 1A and B). Thus, while the rate of RNA synthesis influences the abundance of both premature and mature RNAs, processing and degradation rates impact just on premature and mature forms, respectively.

At the transition between steady states, both the new level of transcript abundance and the speed of the transition (responsiveness) depend on RNA kinetic rates. In the most straightforward case, differential expression - the regulation of the cellular abundance of premature (P) and mature (M) RNA species - derives from changes in the rate of premature RNA synthesis only (k_1_). This implies a change in the amount of nascent RNA for a given gene. While it is often assumed, this should be experimentally confirmed by RNA metabolic labeling before concluding that changes in P or M are transcriptional in nature. In all other cases, differential expression can entail more complex co- and post-transcriptional mechanisms, each governed by a processing (k_2_) and/or degradation (k_3_) rate. Solving the system depicted in Fig. 1B permits to determine the impact one or more kinetic rates can have on the abundance of P and M when modulated over time:

- Constant kinetic rates define steady states where P and M abundances are calculated as k_1_/k_2_ and k_1_/k_3_ ratios, respectively (Fig. 1B,C).
- Modulations in the processing rate k_2_ cause just transient variations in M abundance but permanent alterations in P abundance (Fig. 1D).
- M responsiveness to changes in k_1_ depends on the level of k_3_ (compare Fig. 1E and F) (Friedel et al., 2009; Zeisel et al., 2011), and, more surprisingly, of k_2_ (compare Fig. 1E and G).
- k_1_ and k_3_ can separately generate the same type of M variation if changing in opposite directions (Fig. 1F and H), while only the modulation of k_1_ can affect P (Fig. 1F).
- k_1_ and k_3_ reinforce each other’s modulation of M when changing simultaneously in opposite directions (compare Fig. 1E with I), while neutralize each other’s impact on M if simultaneously adjusted in the same direction (Fig. 1J).

Transient alterations in M induced by a temporary change in k_1_ (Fig. 1K) can be made sharper by a concomitant change in k_3_ (Rabani et al., 2011) (Fig. 1L).

First, these data indicate that measurements of mature RNA are in themselves poorly informative of the real transcriptional state of genes. For example, the detection of a mature RNA is typically taken as indication that the corresponding gene is transcriptionally active. This is not necessarily the case for highly stable RNAs, which might persist long after the gene has become silent. Second, these data illustrate how difficult it is to decipher the mechanism responsible for modulating mature RNAs without determining the corresponding RNA dynamics. For example, the modulation of mature RNA species is typically seen as indication of transcriptional regulation, while it could originate from changes in the dynamics of processing and/or degradation, without any change in the rate of synthesis taking place. Ultimately, these data show the necessity to develop methods for the quantification of RNA kinetic rates, in order to fully disclose the mechanisms behind complex responses in gene expression.

### Experimental and computational pitfalls of RNA metabolic labeling experiments

The system of differential equations introduced in Fig. 1B is undetermined, since it includes two equations and the three unknown kinetic rates. The key to solving this system is to use RNA metabolic labeling with short time pulses, so that the quantification of nascent RNA can be fed into the system as proxy for the synthesis rate (de Pretis et al., 2015). There are two main types of RNA metabolic labeling experiments: one involving the purification of labeled RNA species (Dolken et al., 2008), and the other requiring their chemical derivatization prior to *in silico* identification (Baptista and Dölken, 2018). Both categories of methods are characterized by specific pitfalls, and special care is needed when designing these experiments, particularly when deciding on number of replicates, sequencing coverage, and length of labeled nucleotides pulse (Uvarovskii et al., 2019).

Methods based on the purification of nascent RNA present three main drawbacks: (i) higher costs due to the need to sequence both total and labeled RNA populations; (ii) the need to normalize the signal from the nascent RNA population to that from the total (or pre-existing) RNA population, and (iii) the contamination of labeled with unlabeled (pre-existing) RNA molecules. The need for normalization has been partially addressed by introducing internal standards (Sun et al., 2012), or through computational normalization procedures (de Pretis et al., 2015; Rabani et al., 2014). Rather, the problem of contamination issue is typically acknowledged but left unsolved. To quantify contamination, and to verify whether it varies with the duration of the 4sU pulse, we measured the amount of labeled RNA that can be recovered with pulses of 4sU lasting from 10’ to 2h (Fig. 2A,B). A model based on a constant rate of 4sU incorporation into nascent transcripts did not fit our data, suggesting that the incorporation rate depends on the pulse length (Fig. 2C). Indeed, a model based on an exponential increase of the incorporation rate did fit the data better (log likelihood-ratio test P 2e-27; Fig. 2C,D). A model assuming that the rate of contamination does not depend on the 4sU pulse length, further increased data fitting (P 3.1e-7). Rather, a model in which the contamination increased linearly with pulse length did not improve the fitting any further, and reverted to the constant-contamination hypothesis (P 1; a ≈ 0 in Fig. 2C). Altogether, we determined that 10’-long 4sU pulses, which were often used in these studies (de Pretis et al., 2017; 2015; Fuchs et al., 2014; 2015; Marzi et al., 2016; Michel et al., 2017; Miller et al., 2011; Rabani et al., 2014; 2011; Sabò et al., 2014; Sun et al., 2012), led to 30% of the labeled fraction being originated through contamination of the pre-existing RNA population. Notably, in an independent study in which dendritic cells were subjected to 10’-long 4sU pulses, 30% of the unlabeled RNA was estimated to contaminate the labeled fraction, suggesting that the percentage of labeled RNA being contaminated is even higher (Rabani et al., 2014). Finally, a 30% contamination rate was also reported by (Baptista and Dölken, 2018). As the contamination rate is likely to depend on the cell type, and specific protocol used, it should be reassessed at every experiment, thus further complicating the design of RNA metabolic labeling experiments.

**Figure 2.**
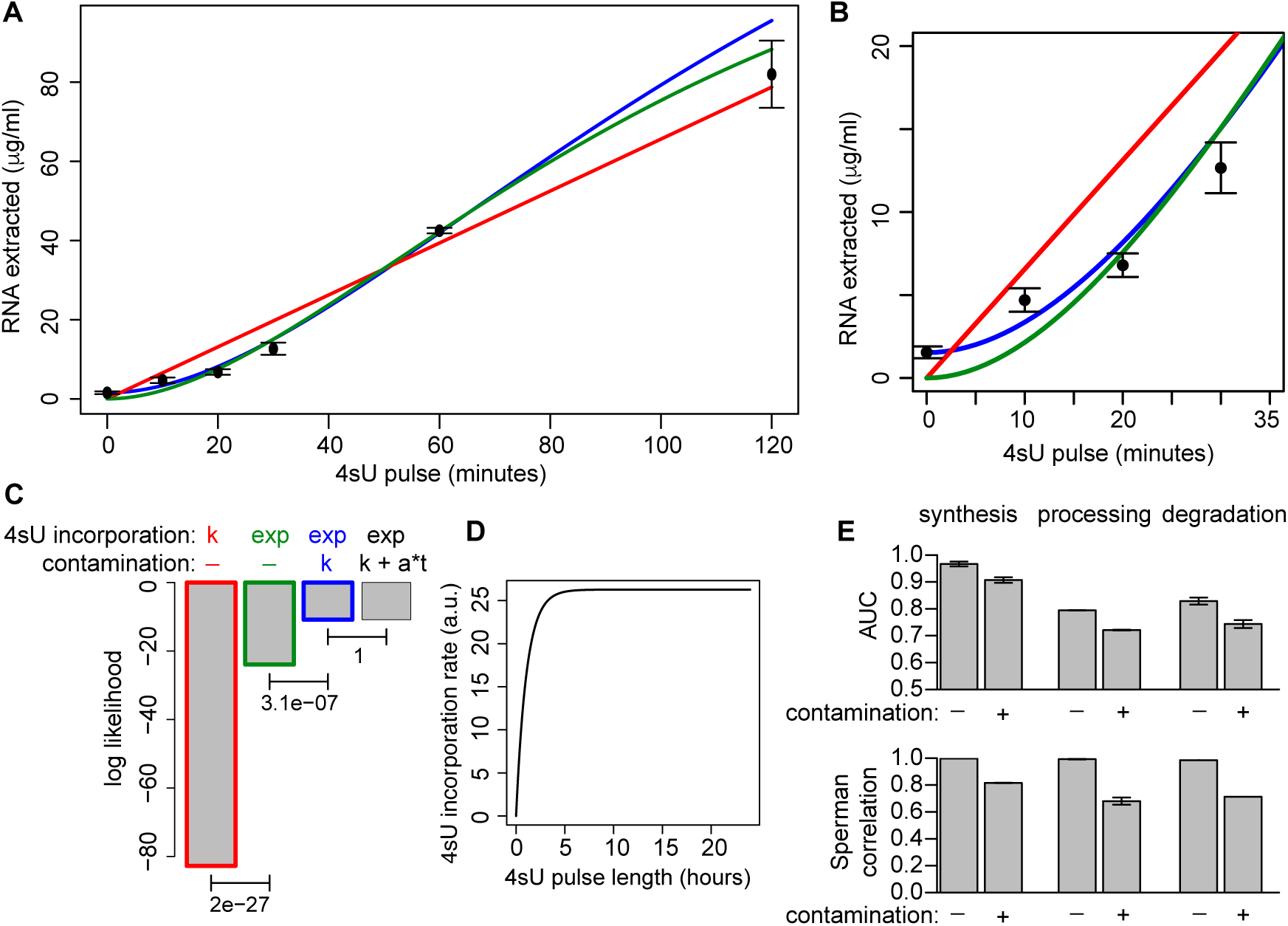
Contamination of 4sU labeled RNA with unlabeled RNA. (**A**) Yield of labeled RNA at 4sU pulses of different length, in 3T9 mouse fibroblast cells. (**B**) A magnification of (A). (**C**) Log likelihood score for the fit of four alternative models, considering constant (k) or exponential (exp) 4sU incorporation rates, combined with a contamination that is absent (-), constant (k), or linear to the 4sU pulse length (k+a*t). P-values of the indicated log likelihood-ratio tests are reported. (**D**) The estimated trend of 4sU incorporation based on the green model in panel (A). (**E**) Changes in ROC Areas Under the Curve (AUC) and in Spearman Correlation following the introduction of 30% contamination in simulated data.

Methods of RNA metabolic labeling that involve chemical derivatization do not rely on the purification of the labeled fraction and therefore do not count normalization and contamination amongst their downsides. However, while able to detect labeled transcripts with excellent specificity, these methods are hampered by low sensitivity and the need for a prolonged pulse time (typically >60’). For example, it has been calculated that 2.4% T>C conversion rates obtained following 24h-long pulses of 4sU in mouse embryonic stem cells (Herzog et al., 2017) permit to identify labeled RNAs at a sensitivity of 23% and 60% for read lengths of 50bp or 150bp, respectively (Neumann et al., 2019). Notably, conversion rates decrease rapidly when the 4sU-pulse length is reduced in order to increase temporal resolution, dropping to 0.5% for a 4h-long pulse (Herzog et al., 2017). A reduced conversion rate is likely to worsen sensitivity. Finally, methods based on RNA metabolic labeling cannot be readily applied to model organisms, mammals (Matsushima et al., 2018) or plants (Sidaway-Lee et al., 2014), *in vivo*.

We recently developed INSPEcT (de Pretis et al., 2015), a Bioconductor package that, together with DRiLL (Rabani et al., 2014), combines analyses on total and nascent transcriptomes to allow, for the first time, quantification of RNA synthesis, processing and degradation rates. Briefly, for each gene, INSPEcT compares eight different models, corresponding to all the possible combinations of each of the three kinetic rates in two alternative analytical forms (constant and impulsive/sigmoid). Each model is plugged within a system of ordinary differential equations (Fig. 1B). The free parameters associated with the rates’ functional forms are optimized to minimize the error when fitting premature and mature RNAs experimental data. Three key aspects of this method have been now updated. First, we have introduced a fully derivative approach able to speed up the execution by 20 fold (Supplementary Fig. 1). Second, model selection has been streamlined, as it now relies on fitting the model in which all rates are variable, avoiding the pair-wise comparison between all nested alternative models. Third, for each kinetic rate, confidence intervals are now determined in order to be exploited for model selection, and to give critical information to the user. As before, INSPEcT is suitable for the analysis of both steady state (Austenaa et al., 2015) and time-course experiments (de Pretis et al., 2017).

INSPEcT’s ability to quantify the kinetic rates absolute values, and to identify genes with variable RNA dynamics, was benchmarked using simulated data that closely reproduced signal and noise of a real dataset (Supplementary Fig. 2 and 3) (de Pretis et al., 2015). However, those data failed to include contamination of the labeled fraction with unlabeled pre-existing RNA, as an important source of bias in RNA metabolic labeling. To measure the importance of contamination, we generated simulated data with and without it. At a 30% contamination level, the correlation with expected rate values, and the Area Under the Curves (AUCs) from ROC analysis, decreased by up to 30% and 12%, respectively (Fig. 2E), indicating that methods based on RNA metabolic labeling are severely affected by contamination of the labeled RNA fraction, and prompting the search for alternative approaches.

### Temporal quantification of RNA dynamics without RNA metabolic labeling

As illustrated in Fig. 1, the modulation of one or more RNA kinetic rates leaves specific marks on the temporal profiles of premature and mature RNAs. Conversely, the temporal quantification of these RNA species should allow the deconvolution of the underlying RNA dynamics. Based on this rationale, we extended INSPEcT to include a novel computational approach able to quantify RNA dynamics using time-course total RNA-seq data, without relying on any RNA metabolic labeling. To keep it simple, INSPEcT+ and INSPEcT-will be used to refer to the application of the INSPEcT package to total and nascent or to just total RNA-seq data, respectively.

Briefly, INSPEcT-follows a 3-step procedure where the ODE system (Fig. 1B) is solved adopting various constraints on the functional shapes of the RNA kinetic rates (Fig. 3A and methods). In the first step (priors estimation), processing (k_2_) and degradation (k_3_) rates are forced to be constant and optimized to reduce the chi-squared error on the mature RNA (M), assuming that premature RNA (P) behaves linearly between the experimental observations. The resulting k_1_ priors, together with P and M, are used in the second step (first guess estimation) to analytically solve the ODE system. This returns k_2_ and k_3_, which are now constant just between experimental time points (constant piecewise). In the last step, M, k_2_ and k_3_ are modeled through a combination of smooth functions (constant / sigmoid / impulsive), minimizing both error and complexity of the model according to the Akaike information criterion (AIC) framework. Finally, k_1_ rates are updated accordingly, and confidence intervals are determined for all kinetic rates. The whole procedure takes approximately 10s per gene per core (Supplementary Fig. 1). See Fig. 3B for an outline of the approaches currently available for the quantification of RNA dynamics, comparing INSPEcT- to INSPEcT+ (de Pretis et al., 2015), DRUID (Lugowski et al., 2018), cDTA (Sun et al., 2012), GRAND-SLAM (Jürges et al., 2018), pulseR (Uvarovskii and Dieterich, 2017), and DRiLL (Rabani et al., 2014).

**Figure 3.**
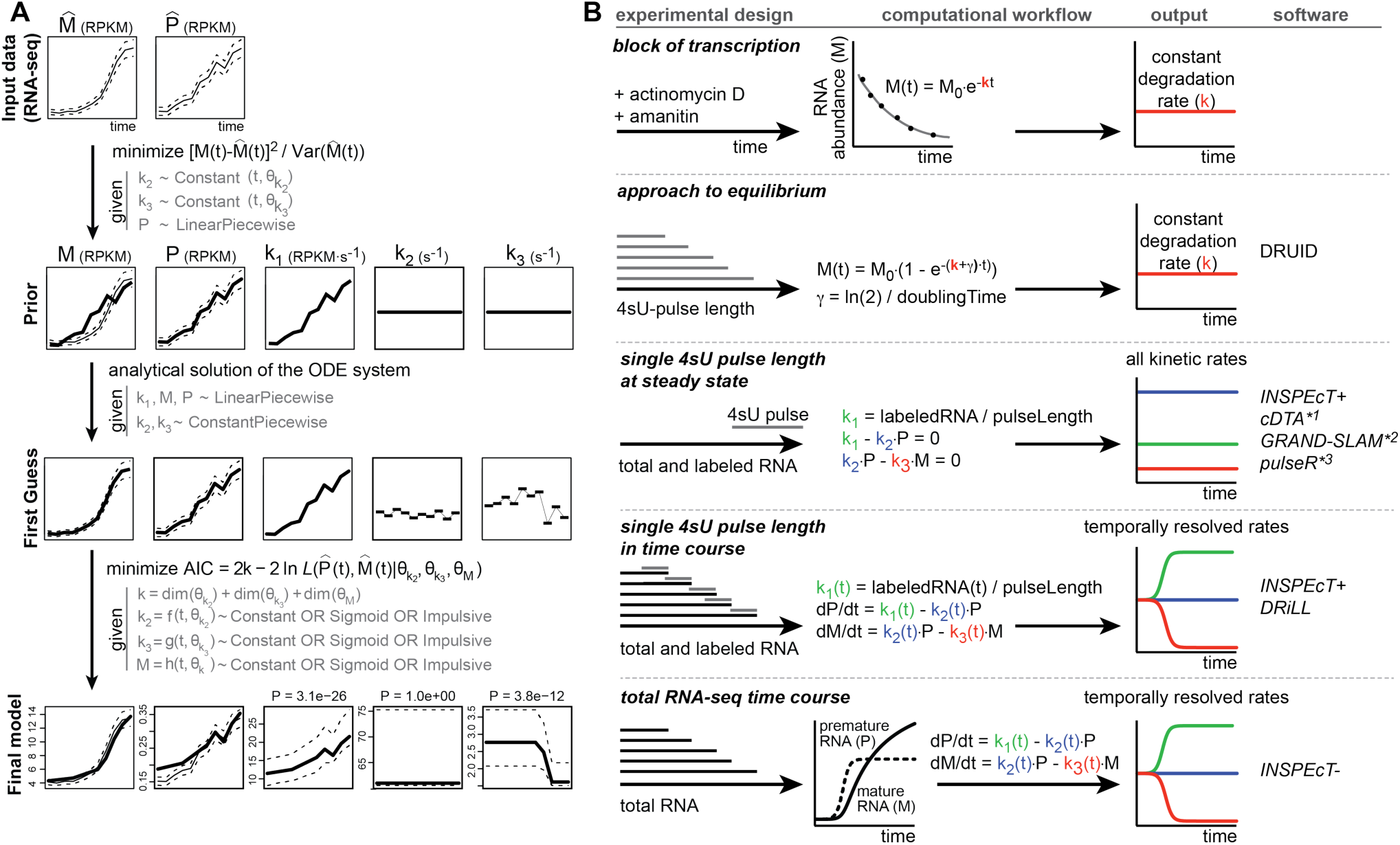
Outline of INSPEcT- and comparison with other approaches. (**A**) INSPEcT-workflow, see text for details; AIC= Akaike information criterion, L= likelihood function **(B)** Experimental design, key features of the computational workflow, output, and software implementation are described for each method. *^1^ it assumes constant processing rates. *^2^ it is based on SLAM-seq data, and does not resolve processing rates. *^3^ it can also analyze time course data but does not temporally resolve the kinetic rates, and does not resolve processing rates.

Fig. 4A shows INSPEcT-output for the H2bc6 gene in 3T9 cells after acute MYC activation, and compares it with the output of INSPEcT+ (de Pretis et al., 2017). This example illustrates the importance of quantifying RNA dynamics quantification: based on classical RNA-seq analyses, the gene in question was not likely to be considered as transcriptionally modulated, since there is little change in total RNA. Instead, H2bc6 is transcriptionally induced more than 2 fold, as the level of premature RNA reveals, while the increase of the mature form is impaired by the rise in post-transcriptional degradation. Additional examples are provided in Supplementary Fig. 4.

We benchmarked the kinetic rates obtained with INSPEcT-in two ways: (i) by comparing their absolute values with those determined by alternative methods, and (ii) by quantifying the ability to detect changing rates in simulated data (Fig. 4B-F).

First we compared the absolute values of all the RNA kinetic rates quantified by INSPEcT-in 3T9 mouse fibroblasts with those provided by INSPEcT+ (de Pretis et al., 2017). Synthesis, processing, and degradation rates had a Spearman correlation of 0.86, 0.61, and 0.69, respectively (Fig. 4B). Second, as the rate of degradation is the only kinetic rate that has been extensively quantified across independent studies, using different experimental and computational approaches, we looked at the RNA half-lives in HEK293 cells. These were quantified by block of transcription experiments and purification of 4sU-labeled RNAs, and the use of DRUID (Lugowski et al., 2018) and INSPEcT+ (Fig 4C). DRUID is based on the approach-to-equilibrium experimental design, which requires to collect various 4sU pulse times (Fig. 3). While being a robust method and a reference in the field, DRUID involves higher costs and more workload, and is poorly suitable for the analysis of changes in decay rates. The mean Spearman correlation between the half-lives quantified by INSPEcT+ and those estimated by DRUID is 0.72 (0.79 when reanalyzing the same data). As it needs just a single 4sU pulse, INSPEcT+ requires only a fraction of the data used by DRUID. Importantly, the 0.72 value above is in line with the correlation between INSPEcT- and INSPEcT+ (Fig. 4B), indicating that absolute quantification of the kinetic rates is possible even in the absence of RNA metabolic labeling data.

**Figure 4.**
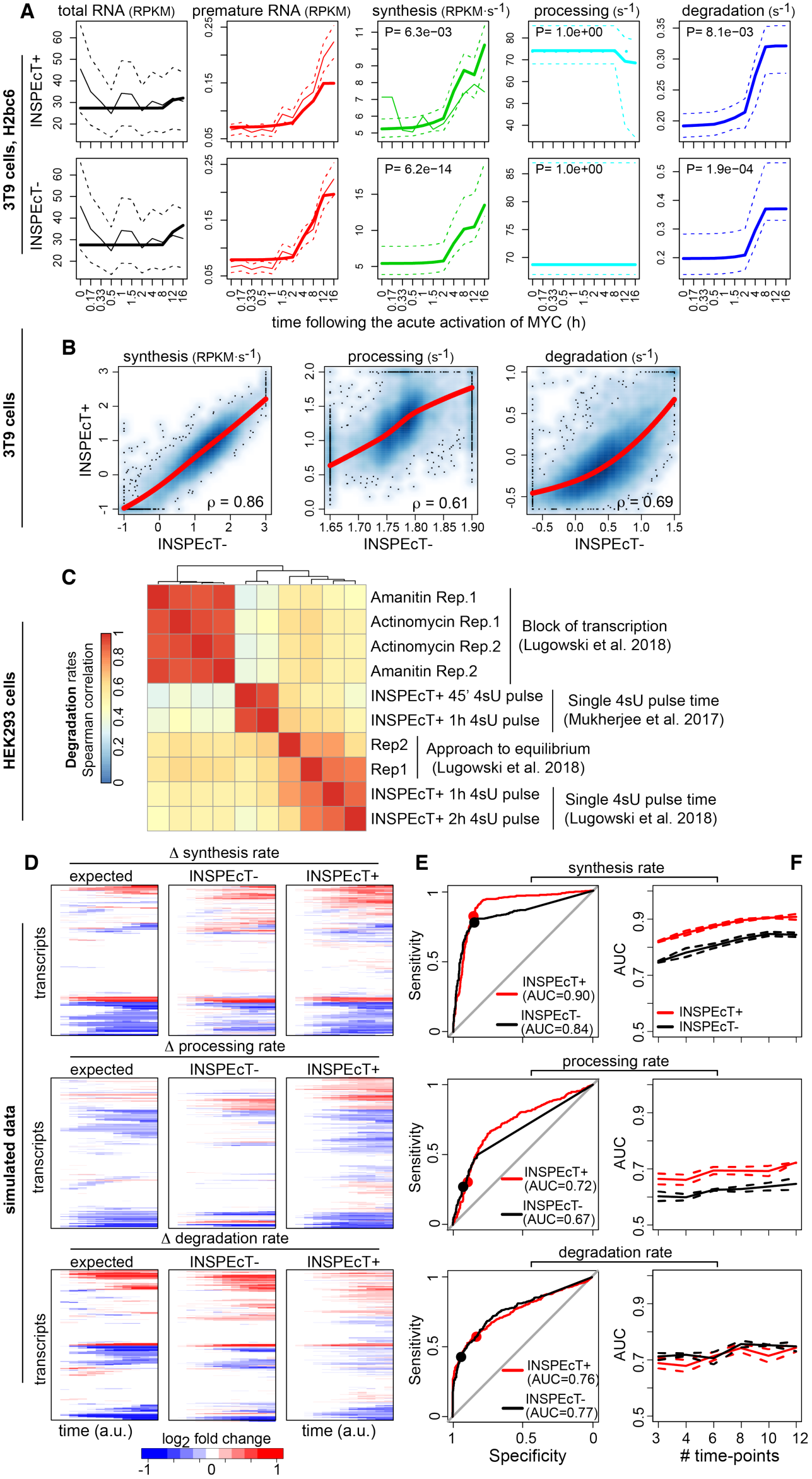
Validation of INSPEcT- kinetic rates. (**A**) INSPEcT-vs INSPEcT+ RNA dynamics for H2bc6. Solid bold lines indicate the model fit; tin solid and dashed lines indicate mean and standard deviation of experimental data for total and premature RNA; dashed lines indicate 95% confidence intervals for the kinetic rates models. (**B**) Comparison of the RNA kinetic rates quantified by INSPEcT+ and INSPEcT-. Regression curves and Spearman’s correlation coefficients are indicated within each panel. (**C**) Heatmap of pairwise Spearman correlations among the half-lives determined through different experimental and computational methods. (**D**) Temporal changes of the RNA kinetic rates for simulated genes, relative to the initial time point (left panels), compared to those quantified through INSPEcT+ (middle) and INSPEcT-(right). (**E**) Sensitivity and specificity in the classification of variable kinetic rates with INSPEcT+ and INSPEcT-, considering 12 time points and three replicates. Dots correspond to P = 0.05. The area under the curve (AUC) is reported within each panel. (**F**) AUCs obtained at increasing number of time points, each including three replicates. AUC’s average and standard deviation are reported based on three simulated datasets.

We then used a simulated dataset of 1000 genes to validate INSPEcT-ability to quantify rates changes. For each gene, both nascent and total gene expression time-course simulated data were included and analyzed using the INSPEcT+ (considering both types of data) and INSPEcT-(considering total RNA data only) approaches. Moreover, the simulated data included matching temporal profiles of RNA kinetic rates, which represented the ground truth for their pattern of modulation (“expected”). INSPEcT-kinetics rates changed over time similarly to INSPEcT+’s, and closely recapitulated the expected response (Fig. 4D). The ability of our procedure of model selection to correctly classify variable rates was measured with ROC analyses and found to perform well (AUC 0.67-0.84; Fig. 4E). These results were in line with those obtained with the INSPEcT+ approach. In particular, sensitivity and specificity at P = 0.05 corresponded to those obtained with nascent RNA. Finally, a reduction in the number of time points did not affect performance significantly (Fig. 4F). These data suggest that the cost of increasing the number of time points is not necessarily justified by an increase in the performance, regardless of nascent RNA profiling.

A possible problem with the INSPEcT-approach lies in the undetermined nature of the ODE system in Fig. 1B. Indeed, the modulation of mature RNA can be potentially explained by changes in either synthesis or degradation. Analogously, the modulation of premature RNA can be potentially explained by changes in either synthesis or processing rates. While this ambiguity can be solved by profiling nascent RNA, which is a proxy for the rate of synthesis, it remains a potential confounding factor when only total RNA is considered. To quantify the importance of this issue, we repeated the ROC analyses by predicting the change in each rate based on the score of the other rates. Swapping the scores decreased INSPEcT-AUCs close to random levels (0.5; Supplementary Fig. 5), indicating that the information gained for different rates is not interchangeable, and demonstrating that indetermination is not a major issue of our approach. Notably, this analysis revealed that INSPEcT+ is more affected by the indetermination issue (Supplementary Fig. 5). In particular, when nascent RNA is profiled, changes in degradation rates can be attributed by error to synthesis and/or processing rates. This is likely due to contamination of labeled RNA with unlabeled transcripts. Indeed, when using simulated data not affected by contamination, the indetermination of INSPEcT+ is fully resolved (AUCs close to 0.5).

Altogether, these analyses indicated that the rates’ absolute values and their changes over time could be estimated even in the absence of nascent RNA data. In particular, INSPEcT-improved the ability to detect changes in degradation rates.

### Reanalysis of public datasets illustrates the additional information gained with INSPEcT-

We used INSPEcT- to reanalyze four publicly available RNA-seq time-course datasets, corresponding to conditions with varying proportions of transcriptional and post-transcriptional regulation (Fig. 5A,B). The analysis of time course RNA-seq datasets is typically limited to the quantification of absolute and differential gene expression, as depicted in Fig. 5C. Rather, INSPEcT-returned the quantification of the temporal changes in premature RNA and of the RNA kinetic rates (Fig. 5D), markedly extending what can be gained from the original data.

**Figure 5.**
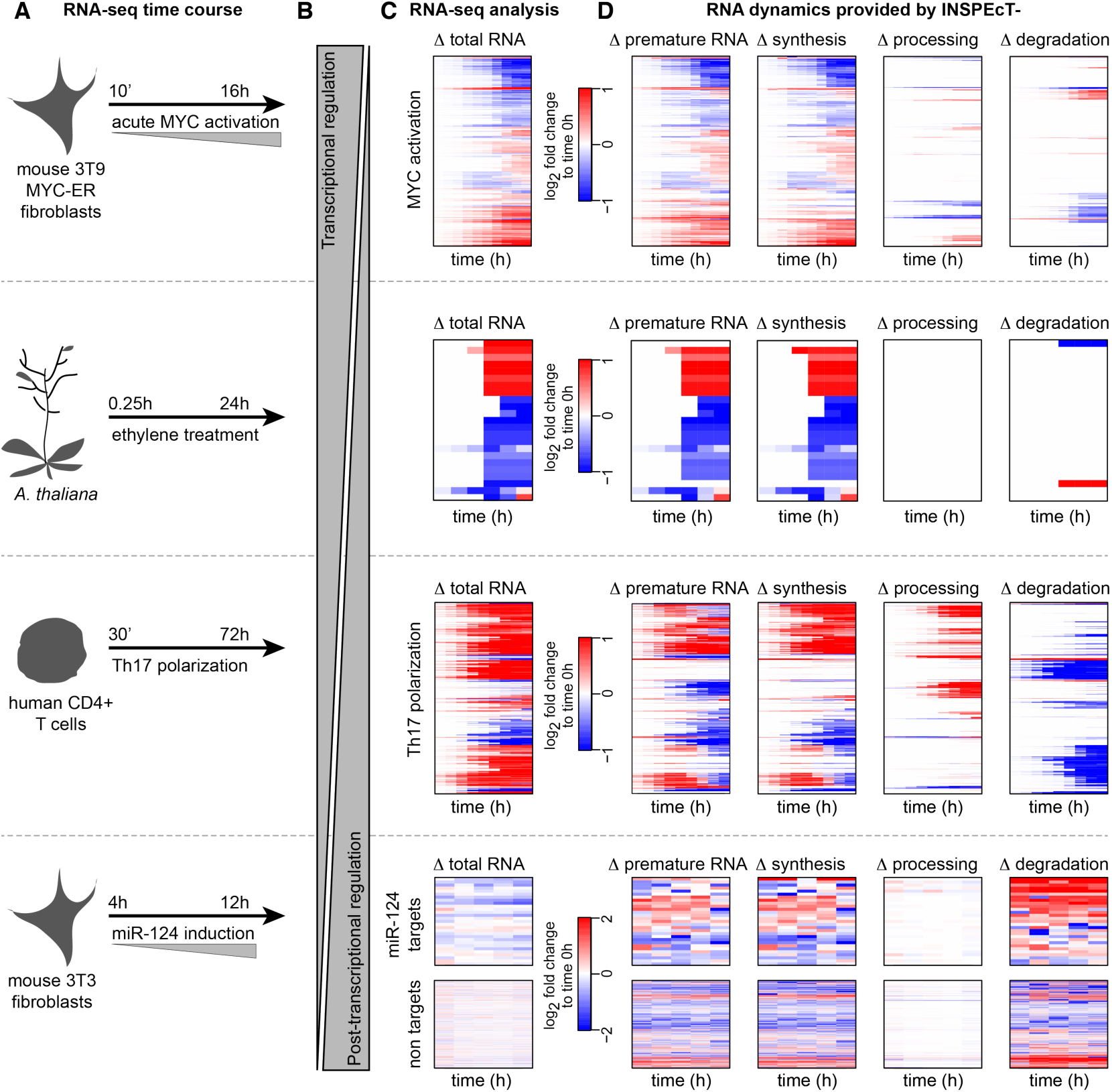
Characterization of time course RNA dynamics: reanalysis of four published datasets with INSPEcT-. (**A**) Experimental design of the considered RNA-seq time courses. (**B**) Expected balance between transcriptional and post-transcriptional responses in the different experiments. (**C**) The typical output of differential RNA-seq analyses: heatmap of differentially expressed genes. (**D**) The additional information gained by reanalyzing those data with INSPEcT-, which includes the gene-level modulation of premature RNAs, and the temporal changes of the kinetic rates of synthesis, processing and degradation. For the miR-124 dataset, reported rates are first-guess estimates, due to lack of replicates in the time course.

First, we focused on the temporal response to MYC acute activation in mouse fibroblasts, which we had recently characterized by profiling both total and nascent RNA (de Pretis et al., 2017). In that study, the integrative analysis of both data types with the INSPEcT+ approach revealed that MYC acts predominantly by modulating the rate of synthesis of its target genes, with an important, albeit less prevalent, impact on processing and degradation involving around one third of targets (de Pretis et al., 2017; Sabò et al., 2014). In agreement with those results, reanalysis with INSPEcT-(which neglects any available nascent RNA data) confirmed that 85% of MYC targets were impacted at the level of their synthesis rate, while 32% of them were affected in either processing or degradation (Fig. 5D).

Second, we quantified for the first time all kinetic rates in plants, focusing on the temporal response to ethylene in *Arabidopsis thaliana* (Chang et al., 2013). Ethylene causes growth inhibition, which is initially independent and then dependent on the EIN3 transcriptional regulator. After 4h EIN3 binding reaches its maximum, leading to a strong transcriptional response (Chang et al., 2013). Indeed, our analysis confirmed that ethylene response is primarily controlled at the transcriptional level (Fig. 5C).

Third, we reanalyzed the total RNA-seq dataset on the temporal polarization of CD4+ T cells with polarizing cytokines from (Tuomela et al., 2016). As expected, in comparison to the one elicited by a master transcription factor of the likes of MYC, the response was more mixed and less dependent on the modulation of synthesis rates: 72% of genes were modulated at the level of their processing and/or degradation rates (Fig. 5D).

Finally, we re-analyzed the total RNA-seq temporal response to the activation of miRNA-124 (Eichhorn et al., 2014). We expected to see a strong and specific post-transcriptional regulation of the miRNA target transcripts and, indeed, these were seen to be primarily controlled at the level of their stability, leading to a reduction in total RNA, while non-target transcripts remained mostly unaffected (Fig. 5C).

Altogether, these analyses illustrate how the quantification of RNA dynamics from total RNA-seq datasets can unveil the underlying mechanisms controlling premature and mature RNA abundances and their variations.

### Temporal quantification of RNA dynamics without assumptions on the functional form

In this study, changes in premature and mature RNA, and in the kinetic rates, are modeled by fitting sigmoid or impulse functions. Sigmoids are the most elementary non-linear functions for modeling a smooth transition between two steady states. Impulse models, which combine an early response followed by an additional transition to a steady state, were previously proposed and successfully used for the modeling of transcriptional responses (Chechik and Koller, 2009; Chechik et al., 2008). Moreover, they were already adopted in the context of RNA dynamics modeling (Rabani et al., 2014; 2011). However, despite their flexibility and broad applicability, these functional forms place a constraint on the modeling, which may poorly adapt to other temporal response patterns, such as oscillatory or more complex responses.

In order to deal with these cases without introducing additional, or overly complicated functions, we implemented a modeling approach based on linear piece-wise functions, available for both INSPEcT+ and -. Briefly, confidence intervals are determined for first guess kinetic rates (Fig. 3A), thus revealing the degree of dissimilarity from a constant model without assuming alternative functional forms. To test this approach, we built datasets including simulated genes modulated by a circadian oscillation of synthesis rates, or by a circadian oscillation of both synthesis and degradation rates opportunely dephased (Fig. 6A). Models returned by both INSPEcT+ and – and obtained by fitting sigmoid or impulse functions, had a poor goodness of fit for the oscillating genes (Fig. 6B). Rather, we found that both approaches can successfully model these circadian oscillatory patterns when agnostic of a priori knowledge of the functional forms (Figure 6C).

**Figure 6.**
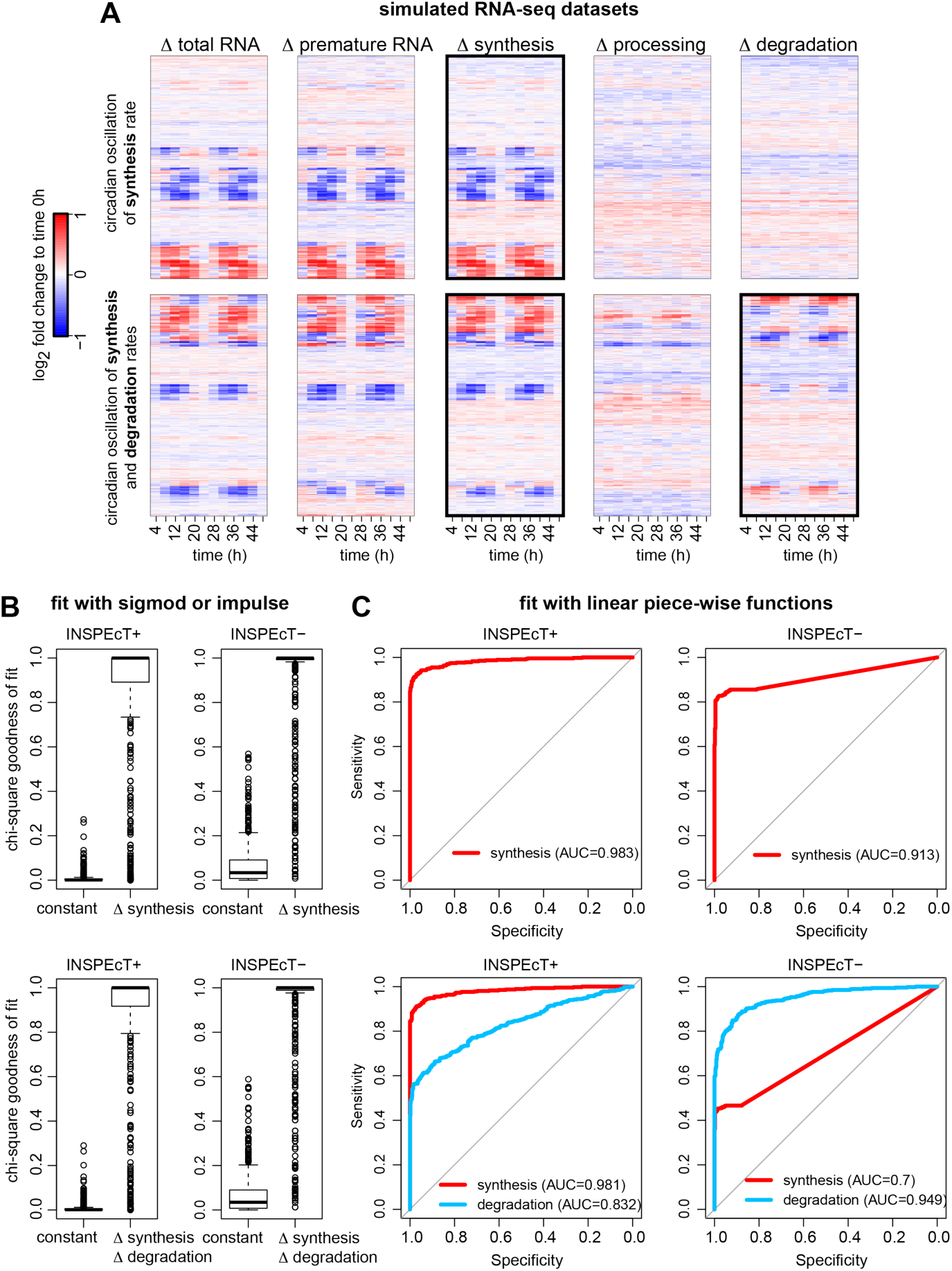
Modeling RNA dynamics without assumptions on the functional form. (**A**) Simulated data composed by 500 constant genes and 500 genes subject to the circadian oscillation of synthesis (top) or synthesis followed by degradation rates (bottom). (**B**) Chi-square goodness of fit of sigmoid or impulse models on the datasets in (A) using INSPEcT+ or INSPEcT-. (**C**) ROC analysis of the classification of synthesis (top) or synthesis and degradation rates (bottom) using INSPEcT+ or – with liner piece-wise functions.

### RNA-dynamics from steady state total RNA-seq data

At steady state and in the absence of nascent RNA profiling, no information on the rate of synthesis is available. However, the ratio of premature to mature RNA abundance is equal to the ratio of degradation to processing rate (k_3_/k_2_, Fig. 1B). While this ratio does not allow the deconvolution of the individual contributions of the two rates, its change over different conditions indicates alterations in post-transcriptional regulation. INSPEcT-uses the ratio of premature to mature RNA species to provide an excellent estimate of the k_3_/k_2_ ratio and its variation (Fig. 7A, B), suggesting that steady state post-transcriptional regulation can be studied even in the absence of RNA metabolic labeling.

**Figure 7.**
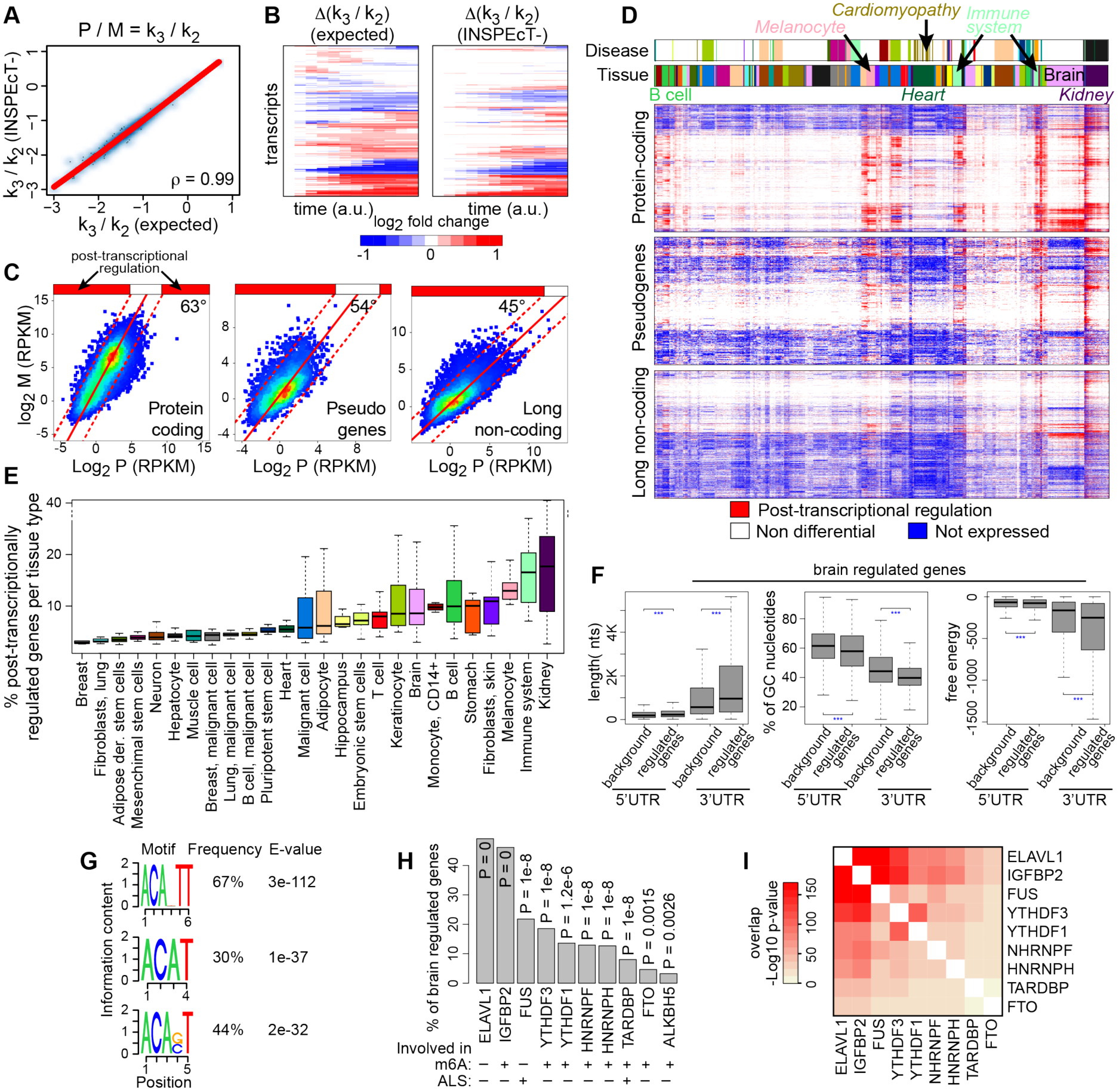
Characterization of steady state RNA dynamics: reanalysis of 620 RNA-seq datasets with INSPEcT-. At steady state the ratio between premature (P) and mature (M) RNA corresponds to the ratio between degradation (k_3_) and processing (k_2_) rates. Absolute values (**A**) and the temporal variation (**B**) of k_3_ / k_2_ ratios determined by INSPEcT-on simulated data were compared to the ground truth (“expected”). (**C**) Median abundances of premature and mature RNAs per gene across 620 RNA-seq datasets for the indicated gene classes. Density scatter plot were fitted with a linear model, whose slope is reported. (**D**) Heatmaps displaying the classification in terms of post-transcriptional regulation for each gene (row) in each sample (column). k_3_ / k_2_ ratios were quantified for each genes in each sample and compared to the global trend depicted in (C). Each gene is either not expressed (blue), not differential (white; ratio between the dashed lines in (C)), or differentially post-transcriptional regulated (red; ratio above the dashed lines). Above the heatmaps, two color bars indicate the tissue-type and disease conditions of each sample. (**E**) Boxplot of the percentage of genes that are post-transcriptionally regulated for the samples associated to each cell type, color matched with (D). (**F**) Distributions of length, %GC, and free energy for 3’ and 5’ UTRs of genes post-transcriptionally regulated in brain, compared to all genes expressed in brain (background). (**G**) Sequence logo of the selected RNA binding protein motifs in the 3’UTR regions of brain regulated genes. (**H**) Frequency and p-values of enrichment for selected motifs of RNA binding proteins found in UTR regions of brain regulated genes. (**I**) Hypergeometric pvalue for the overlap between the genes associated to the factors in (H).

Based on this rationale, we used INSPEcT- to characterize the landscape of human post-transcriptional regulation with an unprecedented breadth, covering 35.000 genes and more than 600 samples. Leveraging natural language processing approaches that we had recently implemented in the Onassis Bioconductor package (Galeota and Pelizzola, 2017), each dataset was assigned to a specific tissue type and disease condition, ultimately covering 26 tissues and 24 diseases. We focused on RNA-seq datasets depleted of ribosomal RNA species and therefore enriched of both premature and mature RNAs. Moreover, we relied on RNA-seq coverage data that had been homogeneously reanalyzed across datasets as a part of the recount2 project (Collado-Torres et al., 2017), thus minimizing potential batch effects due to different analysis pipelines and normalization methods. We found that the amount of premature RNA (P) increases with the abundance of mature RNA (M) following a power-law that depends on the gene type (protein coding, pseudo or long non-coding; Fig. 7C). This suggested that non-coding transcripts have higher proportions of unprocessed RNAs, in agreement with our recent report that this class is characterized by slower processing rates (Mukherjee et al., 2017). Significant deviations from these trends, for each gene class, point to post-transcriptionally regulated genes (Fig. 7C).

Each gene, within each sample, was classified as post-transcriptionally regulated (red in Fig. 7D), non-differential (white), or not expressed (blue). Unsupervised clustering of the heatmap columns resulted in the spontaneous grouping of samples from similar tissues and disease conditions (Fig. 7D, and Supplementary Fig. 6), suggesting that post-transcriptional regulation is coordinated across similar biological conditions. Importantly, the observed sample clustering did not simply arise from gene expression patterns of tissue-specific genes, since it was 30% different from the clustering obtained based on mature RNA (Supplementary Fig. 7). This analysis allowed us to rank samples and genes according to their propensity to be regulated at the post-transcriptional level. On one hand, this revealed that post-transcriptional regulation is particularly common in specific conditions (Fig. 7D, E). On the other hand, this indicated that the three gene classes were markedly different in terms of post-transcriptional regulation, with protein coding and pseudo genes being regulated more frequently than non-coding ones (Fig. 7D). Finally, we analyzed the function of the 1000 protein-coding genes with the lowest frequency of post-transcriptional regulation, and found them to be associated with basic cellular processes such as protein folding, organelle organization and metabolic processes. On the contrary, the 1000 genes with the highest frequency were enriched in miRNA targets, and were found to be related to more specific cellular processes, including various diseases, B-cell activation, autoimmune response, differentiation and morphology.

We analyzed more closely the functionality of the genes undergoing post-transcriptional regulation under specific conditions. Genes altered in T-cell samples were associated with the regulation of T-cell number and proliferation and with immunodeficiency, and often targeted by the E2F1 transcription factor, the master regulator in T-cell proliferation (Zhu et al., 2001). Genes altered in heart samples were associated with cardiac hypertrophy, abnormal contractility and cardiomyopathy. Indeed, a subset of these samples could be associated with the cardiomyopathy disease (Fig. 7D). Focusing on the RNAs regulated in brain, the corresponding genes were associated with several diseases including glioma, autism, neoplasm of the nervous system, and with biological processes such as hormone secretion and synaptic transmission. Compared to genes expressed in the brain, the 3’ and 5’UTR regions in the subset of the regulated transcripts are longer, have a lower percentage of CGs, and lower free energy, (Fig. 7F), indicating a higher likelihood of harboring regulatory motifs. In particular, their 3’UTRs are enriched in motifs containing the ACA sequence. In mammals, ACA is where the majority of N6-methyladenosines (m6A) occur, m6A being the most abundant RNA modification and an important determinant of post-transcriptional regulation (Linder et al., 2015; Roundtree et al., 2017). We used AURA (Dassi et al., 2012; 2014) to search for motifs of RNA binding proteins within these UTR regions (Fig. 7H). Among the enriched motifs we found those for ELAVL1, also known as HuR, an important regulator of transcripts stability (Mukherjee et al., 2011). In addition, we identified motifs for several m6A readers and erasers (Edupuganti et al., 2017). Finally, we identified the binding proteins FUS and TARDBP, important factors in Amyotrophic Lateral Sclerosis (ALS) (Paez-Colasante et al., 2015). Notably, the genes associated to the motifs of these factors have a marked overlap (Fig. 7I). For example, >92% of the genes containing the FUS motif in their 3’UTR, and >78% of those containing the TARDBP motif, also contain the ELAVL1 and IGFBP2 motifs (P < 2e-41). Collectively, these data confirm that m6A-directed post-transcriptional regulation is pervasive in brain (Yoon et al., 2018), and potentially relevant for ALS. Finally, this analysis provided sets of candidate regulated genes, and RNA-binding proteins that could be responsible for their atypical dynamics of expression.

Altogether, these results illustrated the type and range of information that INSPEcT-is able to provide from the study of RNA dynamics when individual conditions are compared in the absence of nascent RNA data.

### The impact of different RNA-seq protocols

The measurement of the abundance of both premature and mature RNA is pivotal in all the approaches that aim to quantify RNA dynamics, with or without nascent RNA. Premature RNA is typically quantified by intronic RNA-seq signals, while the abundance of mature transcripts is obtained by subtracting intronic from exonic signals. Numerous studies support the concept that intronic RNA-seq reads are a robust proxy for the abundance of premature RNA and the rate of RNA production (Ameur et al., 2011; Gaidatzis et al., 2015; Rabani et al., 2014; 2011; Zeisel et al., 2011). In particular, in a recent report (Gaidatzis et al., 2015), a comprehensive analysis was conducted which demonstrated the high correspondence between intronic RNA-seq signals and both nascent and chromatin-associated RNA signals.

Throughout this study, in order to maximize intronic signal, we conservatively decided to take into consideration only total RNA-seq experiments where RNA molecules had not been poly-A selected. However, we found that standard coverage (20M aligned reads) RNA-seq libraries prepared with various protocols, including poly-A selection, are also suitable for these analyses (Supplementary Fig. 8A) (Adiconis et al., 2013). Indeed, Spearman correlations between Ribo-Zero and polyA selection protocols are in the order of 0.85-0.9 for both premature RNAs and their ratios to mature RNAs.

To test INSPEcT- on a polyA-selected RNA-seq dataset, we reanalyzed the temporal response to the induction of RAF. Additional data from the same study revealed that the gene expression response was primarily controlled at the transcriptional level (Uhlitz et al., 2017). Despite the low coverage in the time course of the total RNA-seq samples, INSPEcT-confirmed a modulation in the synthesis rate of 90% of the genes with altered kinetic rates (Supplementary Fig. 8B).

These data and results indicate that there is enough intronic signal available in samples subjected to poly-A selection, despite the depletion of premature RNA species, and that the quantification of premature and mature RNA species is robust to the choice of the RNA-seq protocol, thus broadening the scope of our approaches.

## Discussion

The deconvolution of RNA dynamics from transcriptional genomics data is an emerging field of research, which the development of RNA metabolic labeling has fuelled by enabling the analysis of nascent transcription (Baptista and Dölken, 2018; Dolken et al., 2008; Rabani et al., 2011). We recently developed INSPEcT, a Bioconductor package that, through mathematical modeling of nascent and total RNA-seq datasets, allows the quantification of the kinetic rates governing the RNA life-cycle (de Pretis et al., 2015). We extensively used this tool for the analysis of the RNA dynamics controlling several classes of coding and noncoding transcripts (Austenaa et al., 2015; de Pretis et al., 2017; Marzi et al., 2016; Mukherjee et al., 2017). Aware of the challenges the integrative analysis of nascent and total RNA-seq data poses, we have now expanded the package with INSPEcT- to include the possibility to use total RNA-seq datasets only, without requiring any information on nascent transcripts.

The RNA kinetic rates calculated by INSPEcT-using total RNA-seq experiments, were validated through comparison with those obtained by other methods, and then benchmarked on simulated datasets. In particular, INSPEcT-quantifications of transcripts half-lives were found to be improved compared to INSPEcT+, which is affected by the contamination of unlabeled RNA. By re-analyzing various time-course datasets of total RNA-seq, we proved INSPEcT-’s ability to unravel underlying RNA dynamics and hence provide a deeper understanding of the resulting gene expression programs. INSPEcT-prevents all the additional experimental work required in nascent RNA profiling, and safeguards from a number of pitfalls afflicting RNA metabolic labeling experiments, primarily the difficulty in working with limited RNA amounts and/or tight temporal resolutions, the necessity to normalize the quantification of pre-existing transcripts to that of nascent transcripts, and the contamination of the latter with the former. While at steady state these downsides could be accepted in exchange for the ability to deconvolute all RNA kinetic rates, in time course conditions they might not be justified when considering INSPEcT-straightforwardness. Finally, we also demonstrated that INSPEcT-can unveil RNA-seq dynamics under steady state conditions, by providing the first comprehensive analysis of post-transcriptional regulation using hundreds of publicly available datasets, covering a multitude of tissues and disease conditions. The analysis revealed a signature of brain genes, some of which are involved in ALS, which are potentially post-transcriptionally regulated by m6A RNA modifications.

In conclusion, the characterization of RNA dynamics can uncover the mechanistic details underlying complex transcriptional responses. INSPEcT is a unifying computational tool able to unfold these layers of regulation in most experimental scenarios, independently from the availability of information on nascent transcription, and suitable for both steady state and time courses profiling of total RNA-seq. INSPEcT-provides a new perspective on what knowledge can be gained from total RNA-seq datasets, including those previously published, which can now be used not only for measuring abundance and variation in expression, but also for unveiling the contribution of the different phases in the RNA metabolism. Finally, INSPEcT-is ideal for the identification or prioritization of conditions that are likely to be of high interest to the study of RNA modifications, and of their pivotal role in controlling RNA metabolism (Furlan et al., 2019; Roundtree et al., 2017).

## Author Contributions

M.F., and S.d.P. conceived the method and wrote the software. M.F., S.d.P., and M.P. designed the study. E.G. performed the semantic annotation of the metadata of public RNA-seq experiments. N.d.G. characterized the impact of contamination on RNA metabolic labeling data. E.D. performed the analysis of genes post-transcriptionally regulated in brain. M.F., S.d.P., E.G., M.C., and M.P. interpreted the data. M.F., S.d.P., and M.P. wrote the manuscript.

