## Supplementary Figures for "Genome-wide dynamics of RNA synthesis, processing and degradation without RNA metabolic labeling"

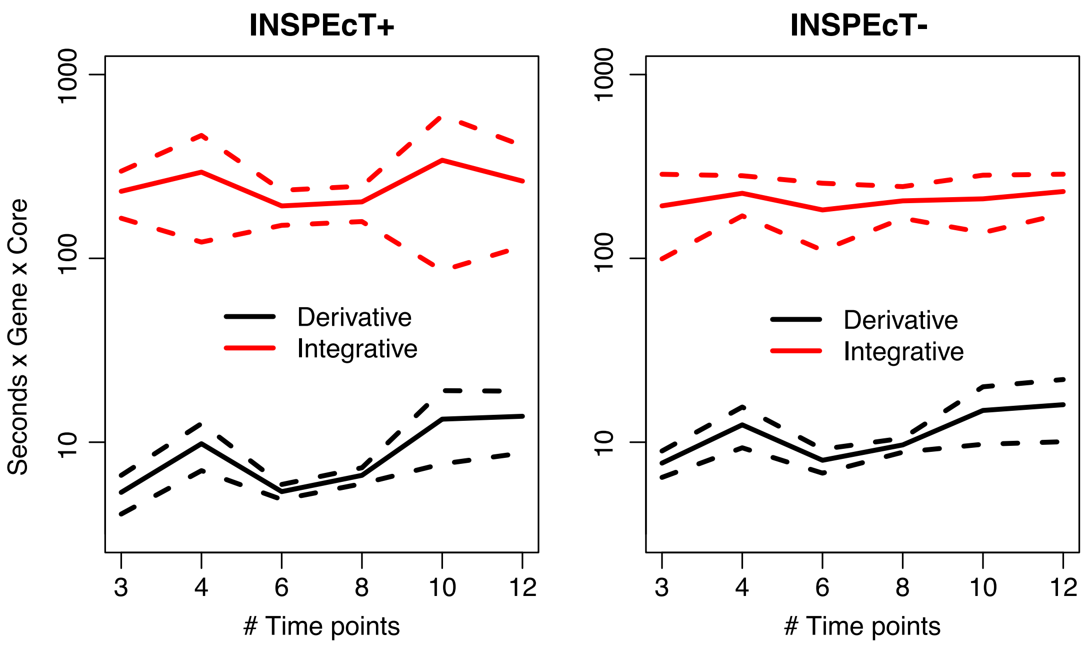


**Supplementary Figure 1 INSPEcT run times per gene.** On the left it is reported the computational times required by INSPEcT+ with derivative and integrative approaches. On the right, the same data are depicted for INSPEcT-. Solid lines represent mean values over ten replicates; dashed lines represent the standard deviation of the mean.

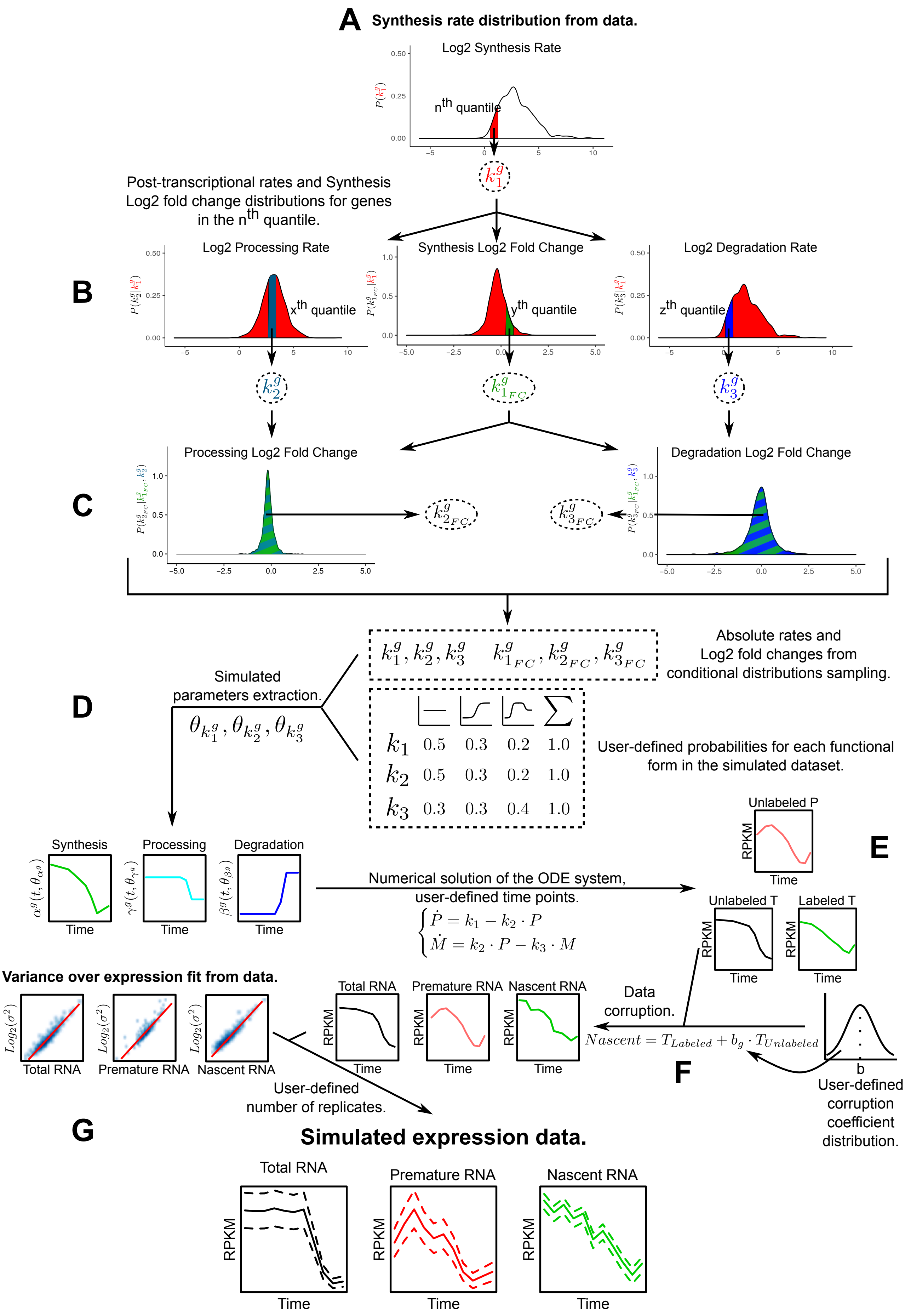

A Synthesis rate distribution from data.
Log2 Synthesis Rate
n^th quantile
Post-transcriptional rates and Synthesis Log2 fold change distributions for genes in the n^th quantile.
B
Log2 Processing Rate
x^th quantile
Synthesis Log2 Fold Change
y^th quantile
Log2 Degradation Rate
z^th quantile
C
Processing Log2 Fold Change
Degradation Log2 Fold Change
D
Simulated parameters extraction.
Absolute rates and Log2 fold changes from conditional distributions sampling.
User-defined probabilities for each functional form in the simulated dataset.
k1 0.5 0.3 0.2 1.0
k2 0.5 0.3 0.2 1.0
k3 0.3 0.3 0.4 1.0
Synthesis
Processing
Degradation
Time
Numerical solution of the ODE system, user-defined time points.
E
Unlabeled P
Unlabeled T
Labeled T
RPKM
Variance over expression fit from data.
Total RNA
Premature RNA
Nascent RNA
Data corruption.
F
b
User-defined corruption coefficient distribution.
User-defined number of replicates.
G
Simulated expression data.

**Supplementary Figure 2 Generation of INSPEcT simulated data.** The procedure is implemented in the *makeSimModel* and *makeSimDataset* INSPEcT methods, and requires the following input: (i) an INSPEcT dataset that includes nascent RNA data, (ii) the number of genes and replicates to be simulated, (iii) a set of time points, (iv) a contamination coefficient together with its dispersion parameter, and (v) the probability of each kinetic rate to be constant, sigmoidal or impulsive.

(**A**) The procedure is performed independently for each gene (g). It starts by sampling the distribution of first guess synthesis rates. (**B**) In order to preserve the correlations among the RNA kinetic rates of the input data, all the genes belonging to the selected quantile are considered. Specifically, the distributions of their processing and degradation rates are determined, together with their synthesis rate fold changes. These distributions are independently sampled to determine these quantities for the gene g (empirical distribution conditioning). (**C**) The procedure is repeated to sample fold changes of processing and degradation rates from their respective conditional distributions. (**D**) Rates temporal profiles are generated based on their magnitude and fold change (the six values returned by the initial part of the procedure), according to the probabilities provided by the user for each functional form. (**E**) Once the simulated kinetic rates are defined, INSPEcT estimates the amount of total and premature RNA within both labeled and unlabeled conditions, by solving the Ordinary Differential Equations (ODE) system at each time point. (**F**) Labeled RNA is optionally corrupted, according to the equation shown in the figure, to simulate its contamination due to unlabeled transcripts. The gene specific corruption coefficient is extracted from a Gaussian distribution whose first and second moments can be defined by the user. For this study, these were set to match 30% contamination rate (based on Fig. 2 results), and 0.72 Spearman correlation between INSPEcT+ and expected degradation rates, thus matching the correlation between INSPEcT+ and DRUID in Figure 4C). (**G**) Finally, to simulate the experimental replicates, we determine the variance expected for each expression datum, based on the power-law global error model implemented in the PLGEM package.

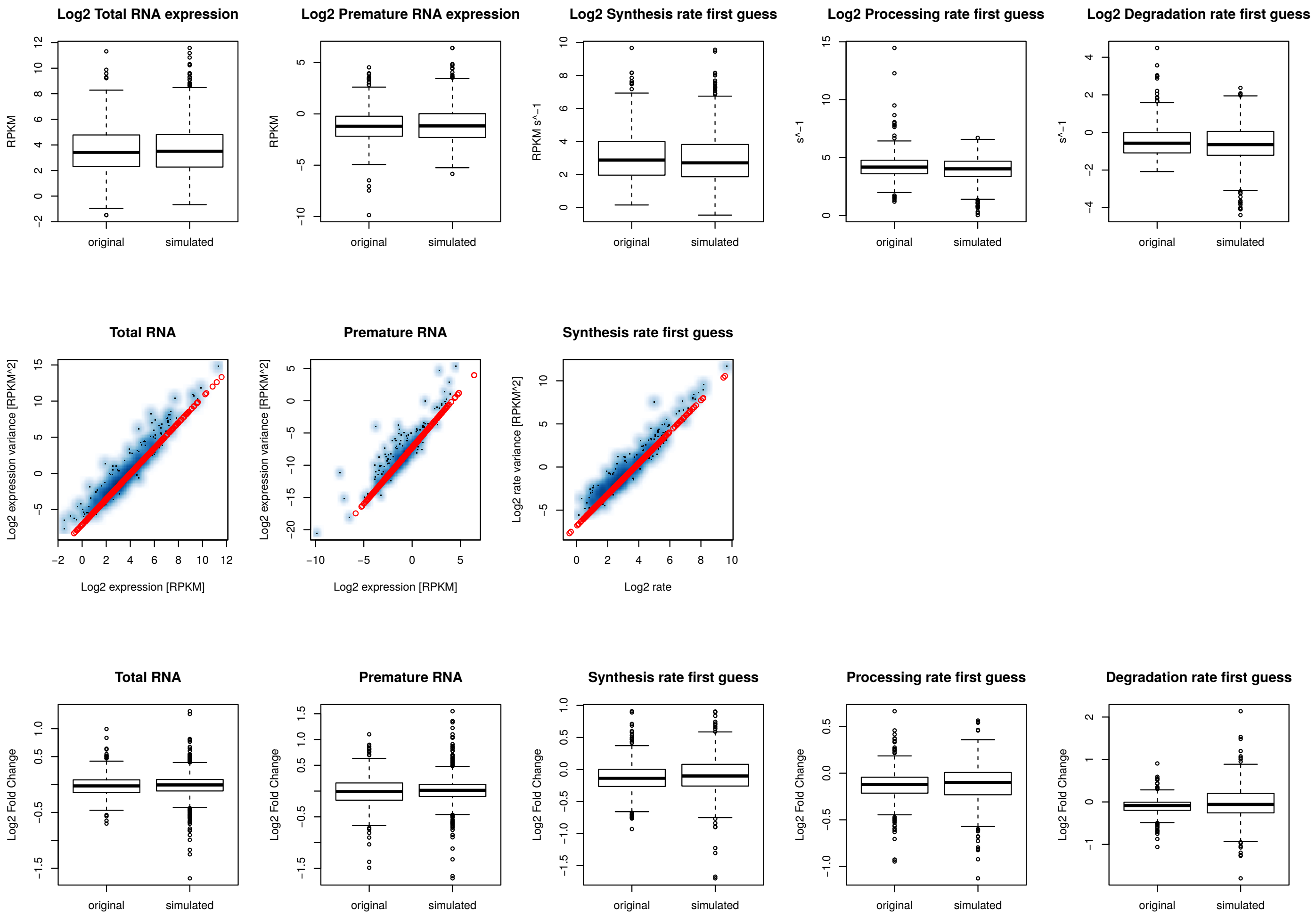


**Supplementary Figure 3 Validation of INSPEcT simulated data.** Absolute levels, variance, and temporal changes of simulated data for 1000 genes (11 time points and 3 replicates) are compared with the original data, a subset of the data from S. de Pretis et al., Genome Research 2017. Simulated data are generated with the *makeSimDataset* and *makeSimModel* functions of INSPEcT, as detailed in the supplementary source code. Boxplots in the first row display Log2 expression data and first guess kinetic rates for original (left) and simulated (right) data. Scatterplots in the second row display the relation between experimental data and their variance in the Log2-Log2 space (density scatter plot in blue); overlaid are the linear fits (determined with the PLGEM package) used to assign a variance to the simulated data. Boxplots in the third row display mean Log2 temporal changes for RNA species concentrations and kinetic rates.

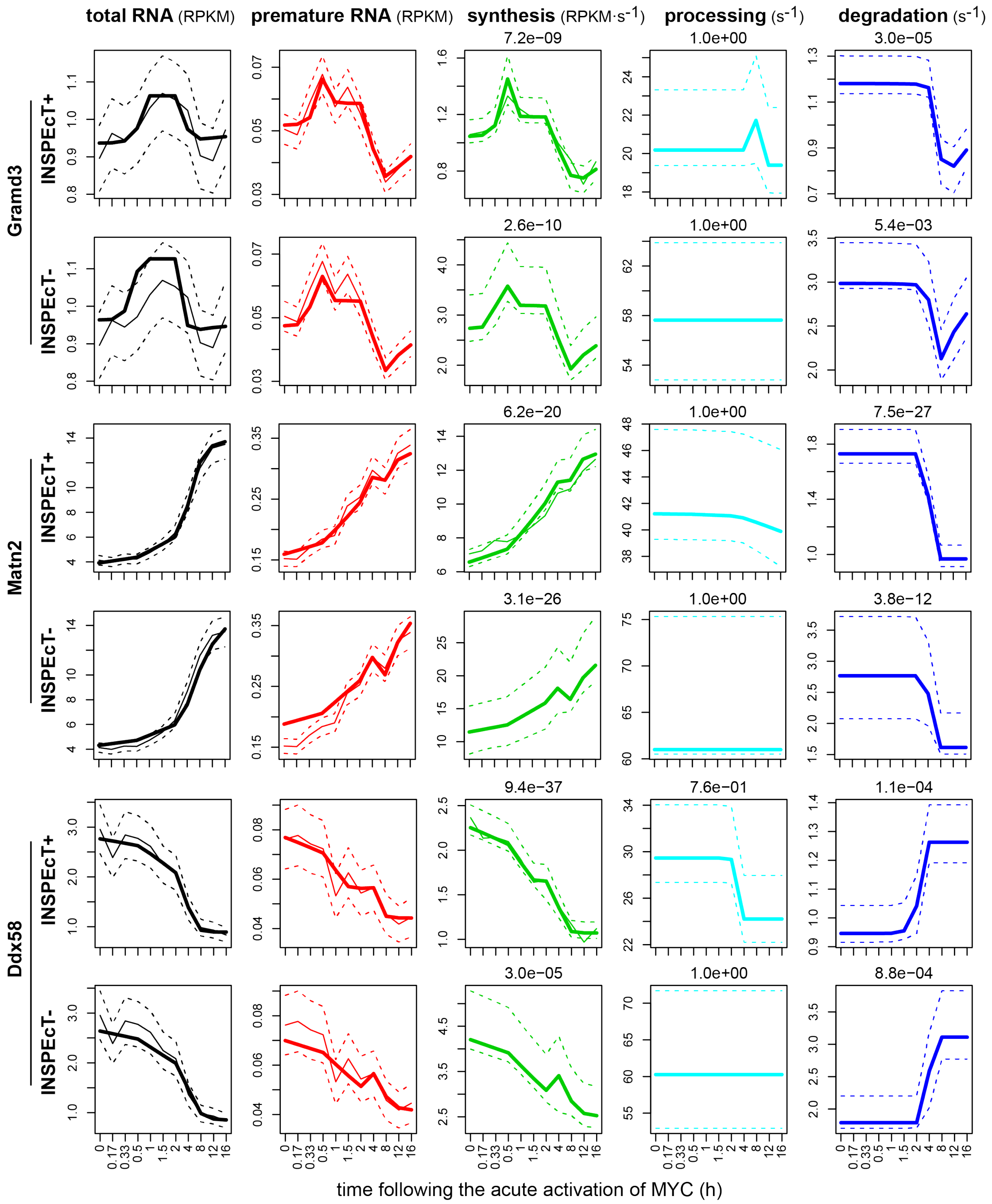


**Supplementary Figure 4 INSPEcT- vs INSPEcT+ RNA dynamics for three genes in 3T9 cells following the acute activation of MYC.** Solid bold lines indicate the model fit; tin solid and dashed lines indicate mean and standard deviation of experimental data for total and premature RNA, respectively; dashed lines indicate 95% confidence intervals for the kinetic rates models. P-values are reported on the top of each kinetic rate indicating the significance of a model in which the rate is varying over time.

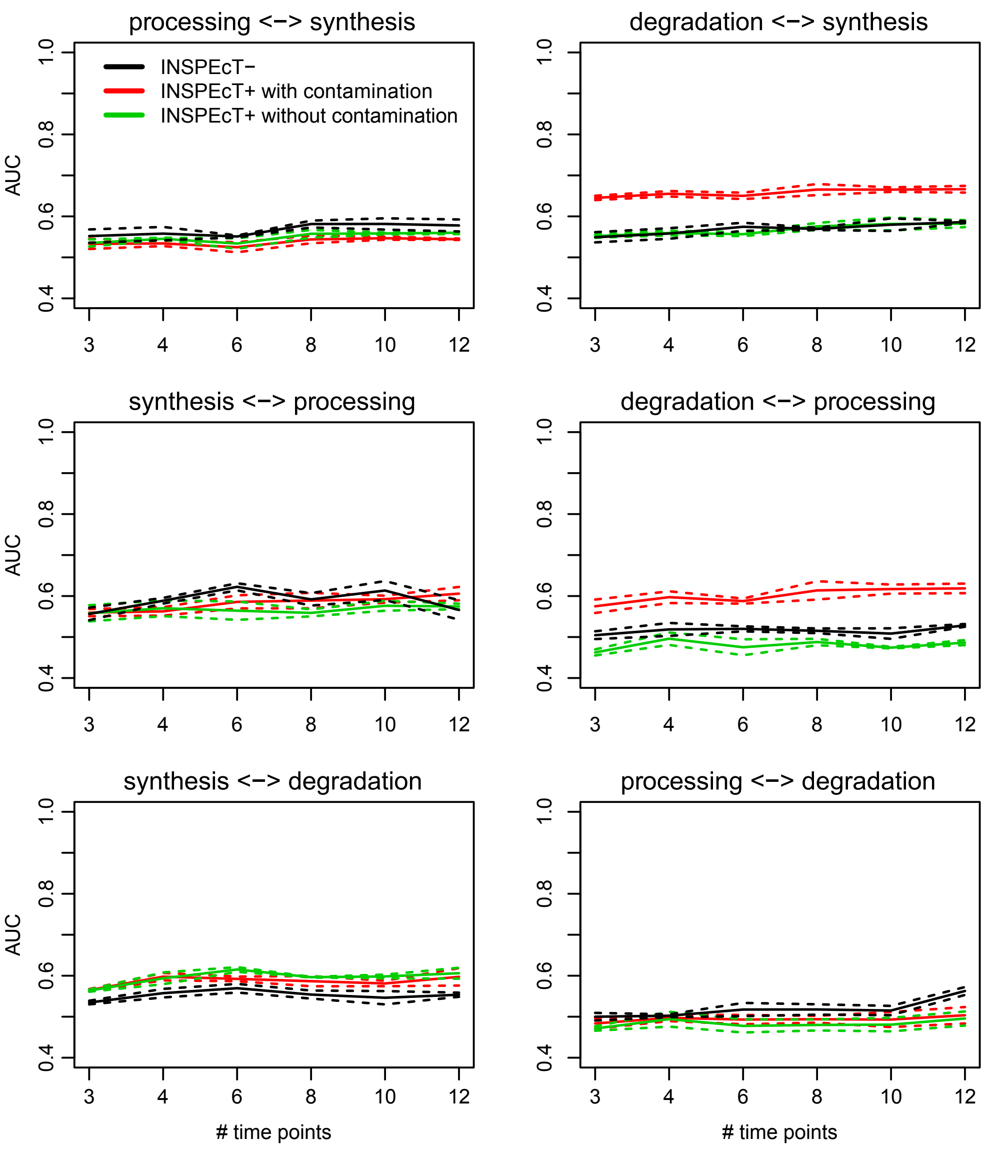


**Supplementary Figure 5 Analysis of the indetermination within INSPEcT- and INSPEcT+.** AUCs obtained by attempting to classify the variability of a given rate (on the left of the arrow in each title) with the estimated variability of another rate (indicated on the right of the arrow in each title). INSPEcT+ (based on simulated data with and without contamination) and INSPEcT– approaches are compared as a function of the number of time points in the simulated dataset. Solid lines are representative of the mean performance over three datasets (1000 simulated genes each); dashed lines show the standard deviation of the mean.

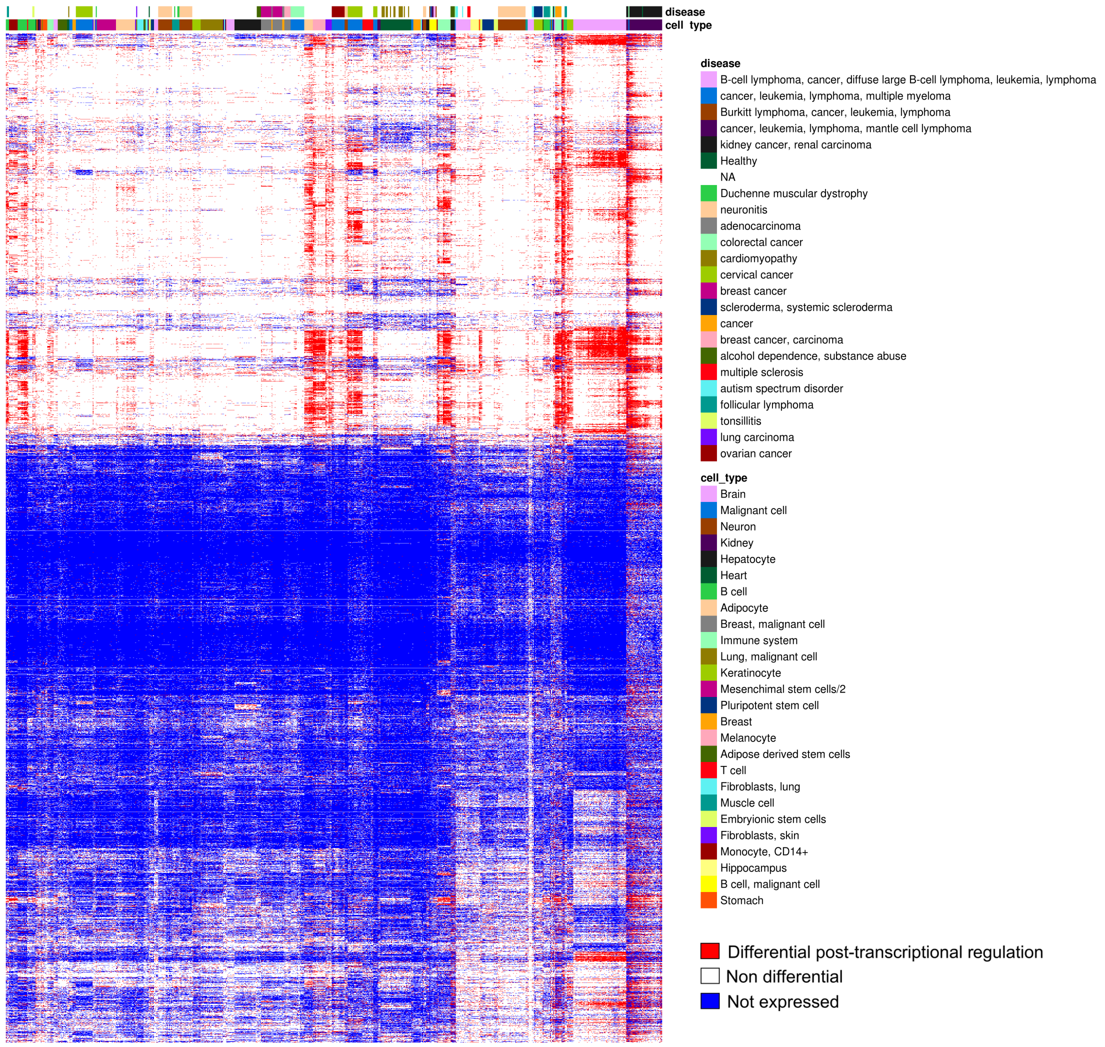


**Supplementary Figure 6 Disease and cell types annotations.** Heatmap displaying the degree of post-transcriptional regulation for each gene (row) in each sample (column). The heatmap combines the three heatmaps displayed in Figure 7D, including the legend of disease and cell type annotations.

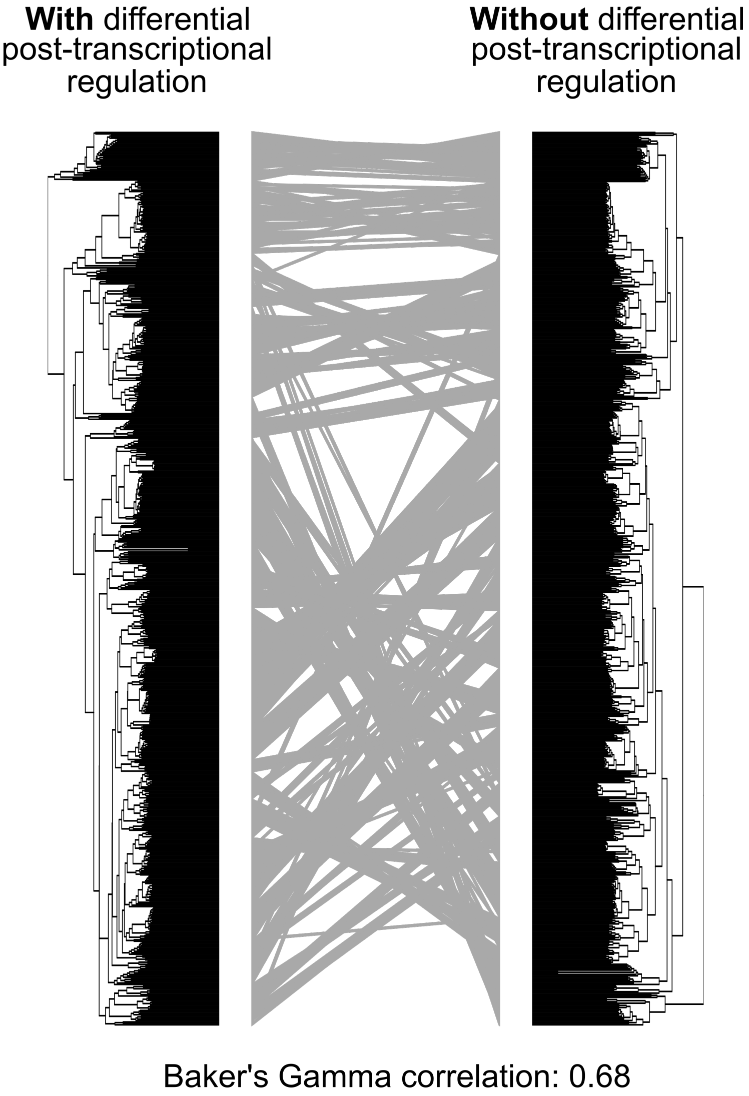


**Supplementary Figure 7 Impact of differential post-transcriptional regulation on the clustering of samples.** Comparison between the hierarchical clustering of samples obtained with (left) and without (right) the information about differential post-transcriptional regulations. In the latter, the underlying data matrix reduces to an object containing binary information: expression or lack of expression of each gene in each sample. Gray edges indicate the repositioning of samples in the two different dendrograms. The Baker's gamma correlation is determined to quantify the similarity between the two dendrograms, denoting >30% difference.

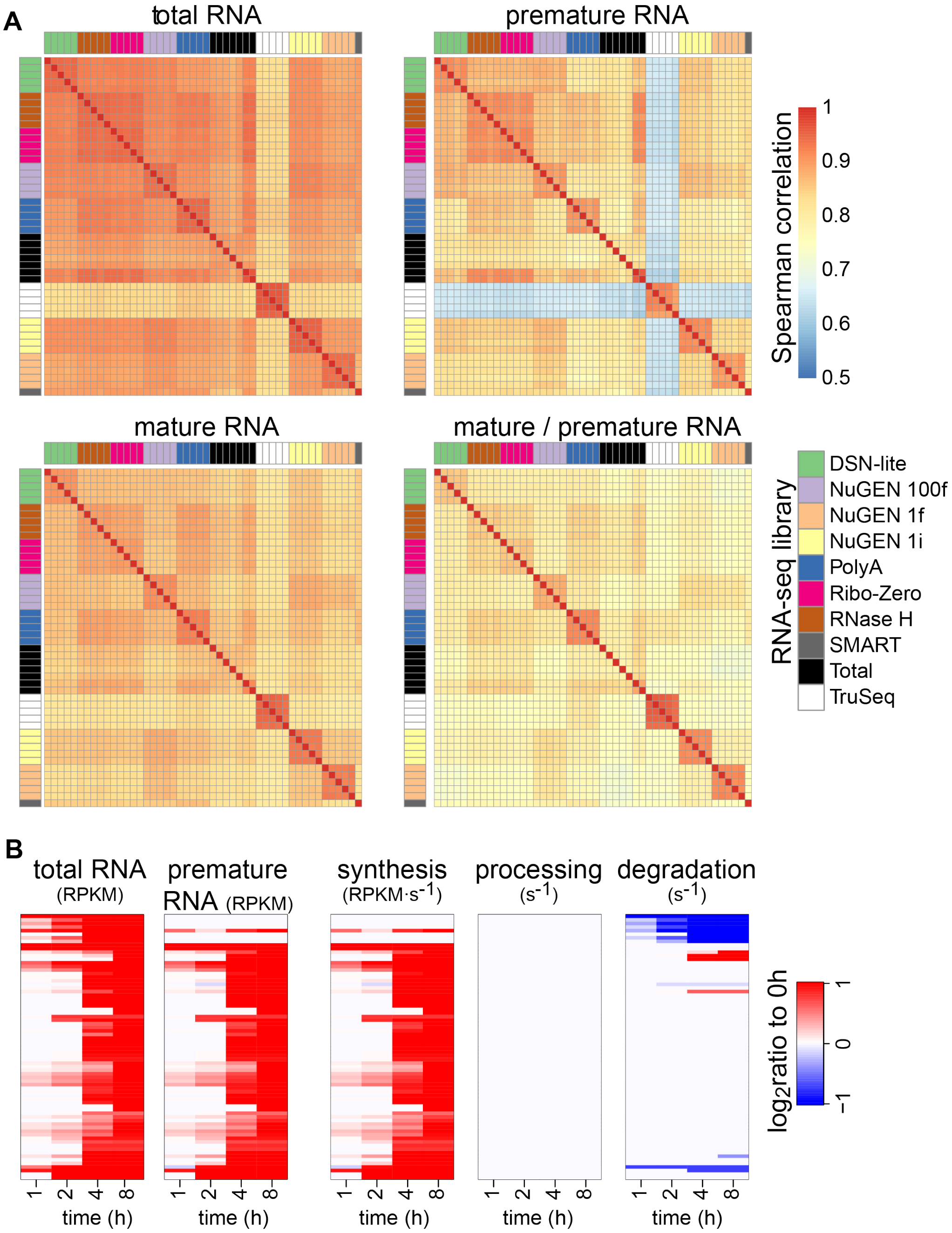


**Supplementary Figure 8 Impact of various RNA-seq library preparation methods on the quantification of RNA species abundance.** (**A**) Spearman correlation of gene-level total, premature and mature RNA abundance through alternative RNA-seq library preparations. (**B**) Temporal changes in total and mature RNA, and in the kinetic rates, following the induction of RAF profiled through RNA-seq of polyA transcripts.
