## Supplementary material for "Genome-wide dynamics of RNA synthesis, processing and degradation without RNA metabolic labeling": Methods

**Quantification of premature and mature RNA expression from total RNA seq data** **2**

**RNA-dynamics from time-course total RNA-seq data (INSPEcT-)** **2**

First guess estimation of the rates 3

Modeling of the rates with sigmoid and impulsive functions 5

Confidence interval estimation 7

Selection of the regulative scenario without assumptions on the functional form 7

**RNA-dynamics from time-course total and nascent RNA-seq data (INSPEcT+)** **8**

First guess estimation of the rates 8

Modeling of the rates with sigmoid and impulsive functions 8

Selection of the regulative scenario without assumptions on the functional form 9

**Identification and annotation of the samples for the steady state analysis** **9**

**RNA-dynamics from steady-state total RNA-seq data (INSPEcT-)** **9**

Definition of gene classes 9

Power-law analysis and identification of post-transcriptionally regulated genes 10

Analysis of the post-transcriptional regulation matrix 11

Characterization of post-transcriptionally regulated genes in brain 12

### Quantification of premature and mature RNA expression from total RNA seq data

The analyses presented in this paper are based on the joint study of premature (P) and mature (M) RNA expression levels. These quantities are evaluated aligning the reads coming from the RNA-seq experiments to the intronic and exonic regions of each gene. Exonic genomic coordinates (chromosome, start site and end site) are easily available from many sources (we used the R packages: TxDb.Athaliana.BioMart.plantsmart28 for our time course analysis on Arabidopsis Thaliana, TxDb.Hsapiens.UCSC.hg19.knownGene for our time course analyses on Homo Sapiens and TxDb.Mmusculus.UCSC.mm9.knownGene for our time course analyses on Mus Musculus: version 3.2.2 for all of them. For our steady state analysis we used the recount2 package function “recount_exons”; annotation GRCh38) while intronic genomic coordinates can be obtained from the gaps between consecutive exons in the same gene. We define the premature RNA expression level as the normalized number of reads (RPKM) aligning to the intronic region. Instead, we evaluate the mature RNA expression level as the difference between the normalized number of reads aligning to the exonic region and P. In case of multiple isoforms, we consider as intronic those portions of the gene which are always spliced.

### RNA-dynamics from time-course total RNA-seq data (INSPEcT-)

We model the dynamics of premature (P) and mature (M) RNA according to the following schema:

$$\emptyset \overset{k_1}{\rightarrow} P \overset{k_2}{\rightarrow} \mathrm{M} \overset{k_3}{\rightarrow} \emptyset$$

Where $k_1$ and $k_2$ are, respectively, the rates of synthesis and processing of the premature RNA, and $k_3$ is the rate of degradation of the mature RNA. Using mass action kinetics, the above system translates in the commonly used system of differential equations:

$$d_tP(t) = k_1 - k_2\, P(t) \quad \textit{[1.1]}$$
$$d_tM(t) = k_2\, P(t) - k_3\, M(t) \quad \textit{[1.2]}$$

From here on, we will adopt the notation $\widehat{X}$ for experimental observations of the variable $X$, while modeled variables will be denoted in their native form ($X$). With nascent RNA, we have an experimental estimation of $\widehat{k_1}$, without, the sole information of total ($\widehat{T}$, exonic quantification) and premature ($\widehat{P}$, intronic quantification) is used to model the RNA dynamics. In particular, we seek for the simplest $k_2 = f(t, \theta_{k_2})$, $k_3 = g(t, \theta_{k_3})$ and $M = h(t, \theta_M)$ functional forms able to recapitulate the experimental premature RNA ($\widehat{P}(t)$) and mature RNA ($\widehat{M}(t) = \widehat{T}(t) - \widehat{P}(t)$) using the equation *[1.1]*. Applying the Akaike Information Criterion (AIC), our problem translates into:

$$min_{f,g,h,\theta_{k_2},\theta_{k_3},\theta_M} AIC = 2k - 2\, ln\, L(\widehat{P}(t), \widehat{M}(t)|\theta_{k_2}, \theta_{k_3}, \theta_M) \quad [1.3]$$

$$\text{where } k = dim(\theta_{k_2}) + dim(\theta_{k_3}) + dim(\theta_M) \quad [1.4]$$

In practical terms, we search for the parameters $\theta_{k_2}, \theta_{k_3}, \theta_M$ that maximize the likelihood ($L$) of several models with different $f, g, h$ used to describe the data and later choose the one that minimize the $AIC$. Following this, we obtain $k_1$ from *[1.1]* as follows:

$$k_1 = d_t P(t) - k_2\, P(t) \text{ where } P(t) = (d_t M(t) + k_3\, M(t))/k_2 \quad [1.5]$$

### First guess estimation of the rates

We compute a first guess estimation of $k_2$ and $k_3$, starting from the assumption that they are constant throughout the time-course, and we use their values for the initialization of the likelihood maximization procedure *[1.3]*, described in the next paragraph. We solve equation *[1.2]* for $k_2$ at $t = 0$ assuming the steady state ($d_t M(0) = 0$), obtaining the following relation with $k_3$ :

$$k_2 = \frac{\widehat{P}(0)}{\widehat{M}(0)} k_3 \quad [1.6]$$

We can rewrite equation *[1.2]* in the form:

$$d_t M(t) = \frac{\widehat{P}(0)}{\widehat{M}(0)} k_3 \cdot P(t) - k_3 \cdot M(t) \quad [1.7]$$

and solve it for all time intervals:

$$\int_{t_i}^{t_i+\delta t_i} d_t\, (M(t) \cdot e^{k_3 \cdot t}) \cdot dt = \frac{\widehat{P}(0)}{\widehat{M}(0)} k_3 \cdot \int_{t_i}^{t_i+\delta t_i} P(t) \cdot e^{k_3 \cdot t} \cdot dt, \quad [1.8]$$

$$\left[M(t) \cdot e^{k_3 \cdot t}\right]_{t_i}^{t_i+\delta t_i} = \frac{\widehat{P}(0)}{\widehat{M}(0)} k_3 \cdot \int_{t_i}^{t_i+\delta t_i} P(t) \cdot e^{k_3 \cdot t} \cdot dt, \quad [1.9]$$

In particular, assuming that $P(t)$ has a linear piecewise behavior between experimental time points:

$$\left[M(t) \cdot e^{k_3 \cdot t}\right]_{t_i}^{t_i+\delta t_i} = \frac{\widehat{P}(0)}{\widehat{M}(0)} k_3 \cdot \int_{t_i}^{t_i+\delta t_i} (a_i + b_i \cdot t) \cdot e^{k_3 \cdot t} \cdot dt \quad [1.10]$$

where $a_i$ and $b_i$ are the intercept and the slope of the linear piecewise function interpolating $\widehat{P}(t_i)$ and $\widehat{P}(t_{i+1})$. Now, also the right hand part of the above equation can be solved and optimized for the $k_3$ that minimizes the chi-square error of the modeled $M(t)$ with the observed one $\widehat{M}(t)$:

$$min_{k_3}\sum_i \left(M(t=t_i) - \widehat{M}(t)\right)^2 :$$

$$\left[M(t)\cdot e^{k_3\cdot t}\right]_{t_i}^{t_i+\delta t_i} = \frac{P(0)}{M(0)}\,k_3\cdot\left[e^{k_3\cdot t}\cdot\ \frac{k_3 a_i + k_3 b_i\cdot t - b_i}{k_3^{\,2}}\right]_{t_i}^{t_i+\delta t_i} \qquad \textit{[1.11]}$$

We then repeat the optimization in *[1.11]* on a two dimensional space ($k_2$ and $k_3$, i.e. releasing the condition in *[1.6]* that binds $k_2$ and $k_3$ ratio with the ratio between $\widehat{P}(0)$ and $\widehat{M}(0)$), by using $\frac{\widehat{P}(0)}{\widehat{M}(0)}k_3$ and $k_3$ computed above as the starting point of the new optimization for $k_2$ and $k_3$, respectively. The synthesis rate is then obtained from equation *[1.1]* and takes the form of a linear piecewise function:

$$k_1(t) = d_tP(t) + k_2\cdot P(t)$$

$$= d_tP(t) + k_2\cdot(a_i\ +\ b_i\cdot t)\ \text{where}\ \ t_i < t < t_i + \delta t_i \qquad \textit{[1.12]}$$

and $d_tP(t)$ is estimated by interpolating the $\widehat{P}(t)$ experimental time-course with cubic splines. In the next step, we release the also the assumption that $k_2$ and $k_3$ are constant during the whole time-course. In fact, we integrate the differential equations *[1.1]* and *[1.2]* assuming that between experimental time points $M(t)$, $P(t)$ and $k_1(t)$ behave linearly (linear piecewise) and that $k_2(t)$ and $k_3(t)$ rates are constant (constant piecewise) and solve them for $k_2^i$ and $k_3^i$, respectively, within each time interval $(t_i, t_i + \delta t_i)$:

$$Solve_{k_2^i}\left[\widehat{P}(t)\cdot e^{k_2^i\cdot t}\right]_{t_i}^{t_i+\delta t_i} = \left[e^{k_2^i\cdot t}\cdot\ \frac{k_2^i m_i + k_2^i q_i\cdot t - q_i}{(k_2^i)^2}\right]_{t_i}^{t_i+\delta t_i} \qquad \textit{[1.13]}$$

$$Solve_{k_3^i}\left[\widehat{T}(t)\cdot e^{k_3^i\cdot t}\right]_{t_i}^{t_i+\delta t_i} = \left[e^{k_3^i\cdot t}\cdot\ \frac{k_3^i a_i + k_3^i b_i\cdot t - b_i}{(k_3^i)^2}\right]_{t_i}^{t_i+\delta t_i} + k_3^i\cdot\left[e^{k_3^i\cdot t}\cdot\ \frac{k_3 m_i + k_3 q_i\cdot t - q_i}{(k_3^i)^2}\right]_{t_i}^{t_i+\delta t_i}$$

This procedure provides a solution of the dataset that works without replicated experiments, with low computational power and that serve as a prior estimate of the rates for the next modeling procedure. As a rule of thumb, this procedure explains variations of $P(t)$ with variations in $k_1(t)$, and variations of $M(t)$ that are not already explained by the variation in $k_1(t)$ with variations mostly at the level of $k_3(t)$ and more rarely in $k_3(t)$.

#### Modeling of the rates with sigmoid and impulsive functions

During the next modeling procedure, we devised two different strategies for the solution of the system where specific functional forms are assigned to the time-courses of rates ($k_1(t)$,

$k_2(t)$ and $k_3(t)$) and concentrations ($P(t)$ and $M(t)$). These functional forms could be constant, sigmoid and impulse functions.

Constant function:

$$f(t, k) = k$$

Sigmoid function:

$$f(t, h_0, h_1, t_0, b) = h_0 + \frac{h_1 - h_0}{1 + e^{-b \cdot (t - t_0)}}$$

Impulse function:

$$f(t, h_0, h_1, h_2, t_0, t_1, b) = \frac{1}{h_1} \cdot \left(h_0 + \frac{h_1 - h_0}{1 + e^{-b \cdot (t - t_0)}}\right) \cdot \left(h_2 + \frac{h_1 - h_2}{1 + e^{b \cdot (t - t_1)}}\right)$$

In one strategy, which we called *derivative*, the former functional forms are assigned (with exceptions shown in the table below) to $k_2(t)$, $k_3(t)$ and $M(t)$. Following this, $k_1(t)$ and $P(t)$ are obtained directly from the system of differential equations *[1.1 - 1.2]* as a function of $k_2(t)$, $k_3(t)$ and $M(t)$, as follows:

$$P(t) = \frac{d_t M(t) + k_3(t)\, M(t)}{k_2(t)}$$

$$k_1 = d_t \frac{d_t M(t) + k_3(t)\, M(t)}{k_2(t)} - k_2(t)\, P(t)$$

In the other strategy, which we called *integrative*, functional forms are assigned to $k_1(t)$, $k_2(t)$ and $k_3(t)$, and $P(t)$ and $M(t)$ are obtained integrating the system of differential equations, as follows:

$$\int_0^t d_x\, (P(x) \cdot e^{k_2 \cdot x}) \cdot dx = \int_0^t k_1(x) \cdot e^{k_2 \cdot x} \cdot dx,$$

$$\int_0^t d_x\, (M(x) \cdot e^{k_3 \cdot x}) \cdot dx = \int_0^t k_2(x) \cdot P(x) \cdot e^{k_3 \cdot x} \cdot dx,$$

While the integrative solution of the system constrains the shape of $k_1(t)$ and give rise to more regularized results, the derivative solution has the advantage to run orders of magnitude faster. For this reason, and considering that often this method applies to the analysis of several thousand genes, we decided to use the derivative solution as the predefined one.

This second modeling procedure is required to reduce the noise associated with the rates estimated in the first guess and to establish the underlying regulative scenario via the comparison of models with different complexities. In particular, per each gene, eight models covering all the possible regulatory scenarios are optimized:

| | $k_1(t)$ | $k_2(t)$ | $k_3(t)$ | *k* | parameters and functional forms of the *derivative* approach | parameters and functional forms of the *integrative* approach |
|---|---|---|---|---|---|---|
| **0** | Constant | Constant | Constant | 3 | M, $k_2$, $k_3$ | $k_1$, $k_2$, $k_3$ |
| **a** | Variable | Constant | Constant | 6/8 | M(t)~Sigmoid/Impulse, $k_2$, $k_3$ | $k_1$(t)~Sigmoid/Impulse, $k_2$, $k_3$ |
| **c** | Constant | Variable | Constant | 6/8 | T(t)~Sigmoid/Impulse, $k_1$, $k_3$ | $k_2$(t)~Sigmoid/Impulse, $k_1$, $k_3$ |
| **b** | Constant | Constant | Variable | 6/8 | T(t)~Sigmoid/Impulse, $k_1$, $k_2$ | $k_3$(t)~Sigmoid/Impulse, $k_1$, $k_2$ |
| **ac** | Variable | Variable | Constant | 9/13 | [M(t), $k_2$(t)]~Sigmoid/Impulse, $k_3$ | [$k_1$(t), $k_2$(t)]~Sigmoid/Impulse, $k_3$ |
| **ab** | Variable | Constant | Variable | 9/13 | [M(t), $k_3$(t)]~Sigmoid/Impulse, $k_2$ | [$k_1$(t), $k_3$(t)]~Sigmoid/Impulse, $k_2$ |
| **bc** | Constant | Variable | Variable | 9/13 | [T(t), $k_3$(t)]~Sigmoid/Impulse, $k_1$ | [$k_2$(t), $k_3$(t)]~Sigmoid/Impulse, $k_1$ |
| **abc** | Variable | Variable | Variable | 12/18 | [M(t), $k_2$(t), $k_3$(t)]~Sigmoid/Impulse | [$k_1$(t), $k_2$(t), $k_3$(t)]~Sigmoid/Impulse |

In the models c, b and bc of the *derivative* approach the imposed functional form are not strictly assigned to the $M(t)$, $k_2(t)$ and $k_3(t)$ in order to constrain the synthesis rate ($k_1(t)$) to be constant, which is a requisite of those models. Per each gene, the best model is chosen based on the minimum Akaike Information Criterion (AIC):

$$AIC = 2k - 2\ ln\ L(\widehat{P}(t), \widehat{M}(t)|\theta_{k_2}, \theta_{k_3}, \theta_M)$$

where $\theta_{k_2}, \theta_{k_3}, \theta_M$ are the parameters assigned to the functional forms of $k_2(t)$, $k_3(t)$, and $M(t)$, respectively, and $k = dim(\theta_{k_2}) + dim(\theta_{k_3}) + dim(\theta_M)$, or the number of parameters of the model, and *L* is the maximum likelihood of the model. Following this, a score of significance associated with the variability of each of the three rates is achieved by the comparison of closest nested models (shown in Methods Figure 1) by the Log-likelihood Ratio (LLR) Test.

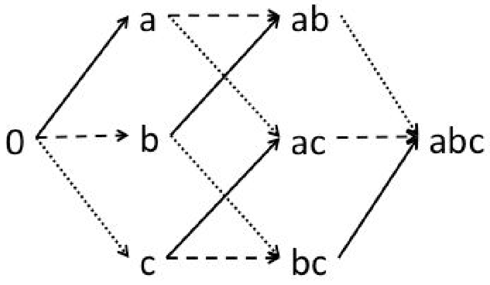


**Methods Figure 1** Explanatory scheme showing how the variability of each rate links different models from the perspective of nesting.

For example, in case the model chosen by AIC is “b”, the significance associated with variable synthesis rate is achieved through the comparison of “b” and “ab”, the significance associated with variable processing rate is achieved through the comparison of “b” and “bc” and the significance associated with variable degradation rate is achieved through the comparison of “0” and “b”. Eventually, a rate is defined variable when the p-value of the LLR test is below the threshold set by the user (by default 0.05) and the final selected model could potentially differ from the model chosen with the lowest AIC.

### Confidence interval estimation

For each parameter ($\Theta opt$) of the selected model, a 95% confidence interval is computed such as, for any $\Theta$ within the interval, the following equation is valid:

$$\frac{logL(\Theta)}{logL(\Theta opt)} < \frac{1}{2}\chi^2_{0.95,1\ d.f.} \quad [3.1]$$

Confidence intervals calculated on the parameters of the model describing each gene are translated into confidence intervals of each rate $k_1(t)$, $k_2(t)$ and $k_3(t)$ by resolving the differential equation system several times, using the optimal parameters for all parameters apart for one parameter at a time, for which the edge of the confidence interval (both the higher and the lower) is used. The ranges of the rates calculated for all these system solutions are used as their confidence intervals.

### Selection of the regulative scenario without assumptions on the functional form

In case the sigmoid or impulse functional forms are not suited to describe the experimental data, such as in oscillatory genes, the rates estimated in the first step of the analysis (see “First guess estimation of the rates”) are used to select the regulative scenario. In this framework, rates are defined piecewise between the experimental time points ($k_1$: linear piecewise, $k_2, k_3$: constant piecewise), therefore not being constrained to any “a-priori” shape. In this case, the time-course of each rate is defined by punctual values corresponding to the time points of experimental observations, and we defined confidence intervals for any of these punctual values using the formula *[3.1]*. We then used these confidence intervals as a measure of the standard deviation and assessed, via the chi-squared goodness of fit test, the probability that the given rate is NOT constant as follows:

$$p(\varkappa^2_{0.95,\,d.f.} \geq D) : D = min_k \sum_t \frac{\left[\widehat{X}(t) - k\right]^2}{(C.I._{min} - C.I._{max})^2} \qquad [4.1]$$

Where the number of degrees of freedom ($d.f.$) is equal to the length of the time-course minus one and $\widehat{X}(t)$ is the time course of the rate under consideration.

### RNA-dynamics from time-course total and nascent RNA-seq data (INSPEcT+)

#### First guess estimation of the rates

In presence of nascent RNA, there is an experimental quantification of the synthesis rate, as follows:

$$\widehat{k_1}(t) = \frac{L_e^t}{t_L} \cdot s_f^t$$

At each experimental time-point $t$, where $L_e^t$ is the exonic quantification in the labeled fraction, $t_L$ is the labelling time and $s_f^t$ is the scaling factor modeled by INSPEcT to normalize the labeled library against the total one (see INSPEcT original paper for details). Given the experimental observations of $\widehat{M}(t)$, $\widehat{P}(t)$ and $\widehat{k_1}(t)$, $k_2(t)$ and $k_3(t)$ constant piecewise rates are obtained by solving the ODE system of equations as described in *[1.13]*.

#### Modeling of the rates with sigmoid and impulsive functions

The next modeling procedure closely resembles the one described in the absence of nascent RNA, with two main differences:

1. The experimental quantification of the synthesis rate $\widehat{k_1}(t)$ is included within the likelihood function $L(\widehat{k_1}(t), \widehat{P}(t), \widehat{M}(t) | \theta_{k_2}, \theta_{k_3}, \theta_M)$
2. Only the parameters of the model named “abc”, the most complex one of the eight described above, are optimized to maximize the likelihood function. Following this, the regulative scenario is selected by estimating confidence intervals of the rates (as in *[3.1]*), and by calculating the probability that each rate is not constant as described in *[4.1]*.

#### Selection of the regulative scenario without assumptions on the functional form

Identical to the procedure used in absence of nascent RNA.

### Identification and annotation of the samples for the steady state analysis

We queried the SRAdb (version 1.40.0) for publicly available non-polyA selected human RNA-seq samples (i.e. *library_strategy* = 'RNA-Seq', *taxon_id* = 9606 and fields *library_construction_protocol*, *design_description* and *sample_attribute* containing at least one among the following keywords: "ribozero", "ribo0", "ribominus" and "ribo-"; and not containing: "cytoplasm", "nascent rna", "poly-a", "polya", "mrna", "ffpe" - i.e. paraffin conserved samples - ; the query was not case sensitive). This procedure selected 3'856 experiments.

The R/Bioconductor package Onassis[2] (version 3.8) was able to annotate 3'724 experiments using either "disease" (https://raw.githubusercontent.com/DiseaseOntology/HumanDiseaseOntology/master/src/ontology/doid-non-classified.obo) or "cell line" (https://raw.githubusercontent.com/obophenotype/cell-ontology/master/cl.obo) ontologies. We removed the following uninformative terms from the annotations obtained with "cell line" ontology: 'cell', 'tissue', 'homo sapiens', 'molecule', 'female organism', 'male organism', 'protein', 'cell line cell', 'chromatin', 'signaling', 'cultured cell', 'multicellular organism', 'compound organ', 'organ', 'nucleus', 'primary cultured cell', 'diploid', 'Bos taurus', 'process', 'chromatin', 'protein', 'size', 'ribosome' , 'organ part', 'time', 'body proper', 'multicellular organism'. Additionally, we set to "Healthy" all the experiments that contained within the fields "sample_attribute" or "experiment_attribute" the following strings: 'healthy', 'disease: none', 'disease: normal', 'disease: presumed normal', 'disease: no ad present', 'disease: no ad evident', 'disease state: normal', 'tissuetype: normal', 'no ad present', 'disease: healthy', 'disease: normal', 'disease: presumed normal', 'disease: none', 'disease: null', 'disease: na', 'disease status: normal', 'tumor: none'. Among the annotated samples, 1140 experiments from 103 projects were found to be part of the recount2 database, which comprehends SRA samples uploaded before February 3, 2016. We selected 1004 experiments from 100 projects annotated in SRAdb with at least 7.5M aligned reads. Finally, data was successfully downloaded from recount2 (version 1.4.5) for 669 experiments from 75 projects. Exonic and intronic quantifications were obtained using the normalized coverage computed by recount2, using the function "coverage_matrix". For experiments with multiple runs associated, mean values were considered. Overall, with this procedure we obtained premature and mature expression quantifications (RPKM) for 35'125 genes in 669 conditions, that were classified in 41 different cell lines and 29 diseases.

### RNA-dynamics from steady-state total RNA-seq data (INSPEcT-)

#### Definition of gene classes

We subdivided the 35'125 genes quantified from recount2 in three classes according to their "gene_type" tag in Gencode annotation (version 25), and we obtained 18'729 protein coding genes (corresponding to the term "protein_coding"), 3'945 pseudogenes (corresponding to the term "pseudogene") and 12'451 long non-coding (corresponding to all other terms).

#### Power-law analysis and identification of post-transcriptionally regulated genes

The power-law, typical of each gene class, was calculated based on the log transformed medians of premature and mature RNA expressions across all samples, and was used as a prototypical combined modulation of transcriptional and post-transcriptional rates upon gene levels regulation. For example, protein coding genes show a greater increase in the levels of mature compared to premature RNA upon up-regulation (Fig. 7C, first box), suggesting a combined action of increased synthesis and processing rates and/or decreased degradation rate (Method Fig. 2). Long non-coding genes, conversely, show an even effect on premature and mature RNA upon gene levels regulation (Fig. 7C, third box), suggesting a major role of synthesis rate on their modulation. Finally, pseudogenes show an intermediate behavior between protein coding and long non-coding genes (Fig. 7C, second box).

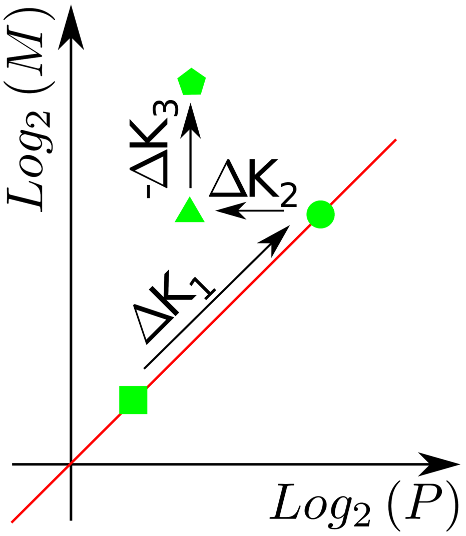


**Methods Figure 2** Effect of each kinetic rate modulation in the Log2(P)-Log2(M) space.

Specifically, the typical power law per each gene class was estimated as the one that included the largest amount of genes within its parallel “acceptance bars”, set 2 units far from the central line in the log2 transformed space of premature and mature RNA medians across samples (Fig 7C).

We hypothesized that significant deviations from these trends points to differentially post-transcriptionally regulated genes (DPR). Specifically, we identified a gene as differentially post-transcriptionally regulated in a specific sample, when resided outside the acceptance bars of the gene class specific power law centered on the median premature and mature values of the gene under consideration (Method Fig. 3).

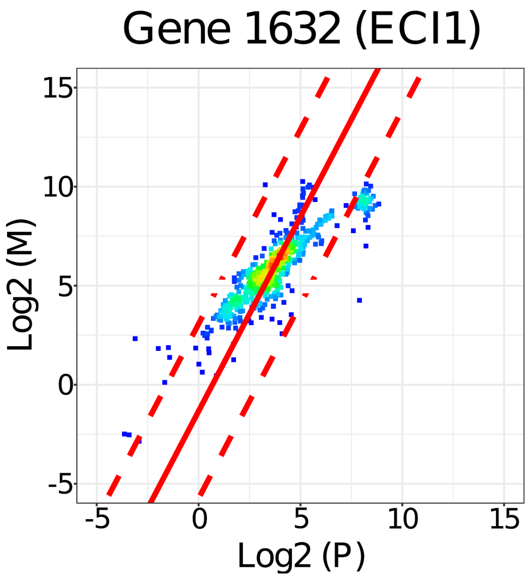


**Methods Figure 3** Log2(P)-Log2(M) plot for a single protein coding gene together with the linear model fitted on the entire gene class. For each point outside the acceptance bars of the model it is associated a RNA-seq sample in which the gene can be classified as differentially post-transcriptionally regulated.

### Analysis of the post-transcriptional regulation matrix

This analysis involved the 26 most represented cell types (620 samples) in order to allow an effective discrimination of the colors used for their identification. For the same desire of clarity , we represented only the genes with at least 310 finite k2/k3 ratios. This choice had almost no consequences on protein coding genes, on the other hand, it changed significantly the graphical impact of the plots for the other two gene classes allowing a less dispersive convey of the information.

We ordered the obtained dataset using hierarchical clustering (using euclidean distance) both on row and columns of the matrix, assigning a value of 0 to non-expressed genes (having at least 1 exonic and 1 intronic read in at least 10 samples), 1 to expressed genes, and 2 to expressed and DPR genes.

We compared the tree obtained from the clustering of the columns with an analogous object produced from a matrix identical to the previous one except for DPR genes which were assigned to 1. In this way, we estimated the amount of information conveyed by this class of entries. To perform the comparison, we used Baker's Gamma function as implemented in the R package dendextend version 1.6[3]. This test resulted in a correlation value of 0.68 (Supp. Fig. 8).

On this dataset, we performed a functional enrichment analysis of the 1'000 genes with the highest number of non DPR samples (not differentially post-transcriptionally regulated genes) and of the 1'000 genes with the highest number of DPR samples (differentially post-transcriptionally regulated genes). The terms of the enrichment where selected imposing simultaneously the same threshold (10^-3) on both the hypergeometric and binomial tests raw p-values trying to compensate for the biases which affect these two metrics. The very same procedure was used to verify enrichments for the most atypically regulated genes for specific groups of experiments identified according to their proximity

after the matrix clustering (hierarchical agglomerative clustering based on the euclidean distance). In this case, we asked for those genes with at least a given percentage of red entries, their number oscillated from some hundreds to some thousands.
compareSteadyNoNascent is the reference function of the INSPEcT package to perform the analysis described above, an example is presented in the vignette.

### Characterization of post-transcriptionally regulated genes in brain

The 5' and 3' UTR sequences of regulated genes were obtained from the UCSC Genome Browser (hg38 assembly, doi: 10.1093/nar/gky1095), along with their predicted secondary structure folding. Differences in length, GC content and free energy between background and regulated genes for each tissue were computed by means of a Wilcoxon test, and plotted with R (https://www.R-project.org/). De novo sequence motifs were searched by means of DREME (doi:10.1093/bioinformatics/btr261), using the sequence of UTRs of the same type from non-regulated genes of the same tissue as the background set, an E-value threshold of 0.05 and considering only motifs on the forward strand. Eventually, to identify known binding sites of regulatory factors in those UTRs, we applied the Regulatory Enrichment tool of the AURA2 database (doi:10.4161/trla.27738) on the regulated genes set of each tissue separately, using an enrichment significance threshold of 0.05 (BH-adjusted p-value).
